# Acetylcholine drives astrocytic JAK2-STAT3 signaling to modulate male-to-female approach behavior

**DOI:** 10.64898/2026.09.02.748883

**Authors:** Tom Lakomy, Camille Falconnier, Kai-Yi Wang, Tony Barbay, Valentin Grelot, Sydney Barthelemot, Martine Guillermier, Anaïs Chekroun, Caroline Jan, Elisa Degl’Innocenti, Maria-Angeles Carrillo de Sauvage, Rémi Bos, Alexandre Charlet, Carole Escartin, Lucile Benhaim

## Abstract

Astrocytes are essential regulators of neural circuits and behavior, sensing synaptic activity and modulating plasticity through diverse intracellular mechanisms. However, the contribution of astrocyte transcription factor-based cascades, linking external stimuli to long-term transcriptional programs, is currently misunderstood. Here we show that the JAK2-STAT3 signaling plays a critical role in male-to-female social behavior. Hence, in male mice, exposure to estrus females rapidly and selectively induces STAT3 signaling in ventral hippocampal astrocytes. Surprisingly, this induction is not driven through canonical cytokine signaling but by septo-hippocampal cholinergic inputs acting through α7 nicotinic acetylcholine receptors (α7nAChR). Astrocytic α7nAChR-JAK2-STAT3 signaling in turn modulates vCA1 pyramidal neuron activity and is required for context-appropriate male approach behavior. These findings reveal a previously unsuspected cellular pathway integrating neuromodulatory inputs and transcription factor-based signaling to shape male behavioral response depending on female receptivity.

## Introduction

Astrocytes are now widely recognized as essential regulators of neuronal circuits, capable of sensing synaptic activity and modulating plasticity ^1–3^. This finely-tuned astrocyte-neuron dialogue shapes a broad spectrum of behaviors across distinct brain circuits ^4–7^. As opposed to astrocytic calcium (Ca2+) signaling, transcription factor (TF)-based pathways - which links external stimuli to long-term transcriptional programs - have been largely overlooked, partly as they operate within longer timescales than Ca2+ transients. Yet mounting evidence indicates that various TF-based signaling are critical for astrocyte regulation of neural circuits underlying adaptive behaviors ^8–10^.

A prime example of such TF-based signaling is the signal transducer and activator of transcription (STAT) family. These proteins are latent cytoplasmic TF that, upon activation, rapidly induce the transcription of quiescent or low-activity genes ^11^. This ubiquitous pathway is classically associated with immune and inflammatory responses, as it is potently activated by cytokines ^12^. In mature astrocytes, the JAK-STAT signaling has been extensively examined in pathological settings, where it drives astrocyte reactivity ^13–19^. However, whether and how astrocytic JAK-STAT signaling contributes to physiological astrocyte-neuron communication remains unknown.

Evidence from both humans and rodents suggests that immune dysregulation impacts various aspects of social interactions ^20^. Interestingly, the link between immune function and social behavior appears to pre-exist, in the absence of an immune challenge. Indeed, a previous study showed that interferon-γ signaling via JAK-STAT in inhibitory neurons-but not microglia-is required for normal sociability in mice, illustrating the increasing recognition of neuro-immune mechanisms are as regulators of social behavior ^21^. Yet, the involvement of astrocytes remains unknown.

Based on this, we hypothesized that the JAK-STAT signaling, a neuroinflammation-associated TF-based pathway, might play a role in neural circuits mediating social behavioral adaption in the adult mouse brain, by coordinating the expression of responsive genes in astrocytes. We focused on male-female interactions, a salient and ethologically relevant stimulation that demands rapid adaptation of male behavior according to female reproductive state. We show that STAT3 - the most abundant STAT isoform in mouse astrocytes - is selectively induced in male mice following exposure to females, in an estrous cycle-dependent manner. Strikingly, we find that astrocytic JAK2-STAT3 signaling is recruited downstream of cholinergic inputs, but not canonical cytokine signaling, and is required for adaptive approach behavior in males. This unconventional pathway gates neuromodulation and TF-based signaling to fine-tune male behavioral responses to female receptivity.

## Results

### Exposure to receptive female induces astrocytic STAT3 signaling in the vHPC of male mice

Since the JAK-STAT signaling is ubiquitous ^11^, we first assessed the expression levels and astrocyte enrichment of several *stat* isoforms, using available RNA sequencing datasets ^22^ (**Extended Data Fig.1a**). Among 7 *stat* isoforms, *Stat3* displayed the highest expression levels in astrocytes (**Extended Data Fig.1b**). We also found that astrocyte *Stat3* enrichment was the highest in the hippocampus (HPC) as compared to the other 10 brain regions (**Extended Data Fig.1c**). We used the astrocyte reporter mouse line Aldh1L1-GFP and compared STAT3 expression pattern between limbic brain regions associated with social behavior ^23^ (**Figure 1a**). Consistent with results obtained at the RNA level, STAT3 protein was predominantly expressed in Aldh1L1-GFP^+^ astrocytes (> 80%) only in the HPC but not in other limbic regions. Furthermore, the numbers of STAT3^-^ astrocytes and of STAT3^+^ non-astrocyte cells were higher in these brain areas than in the HPC (**Figure 1b**).

**Figure 1.**
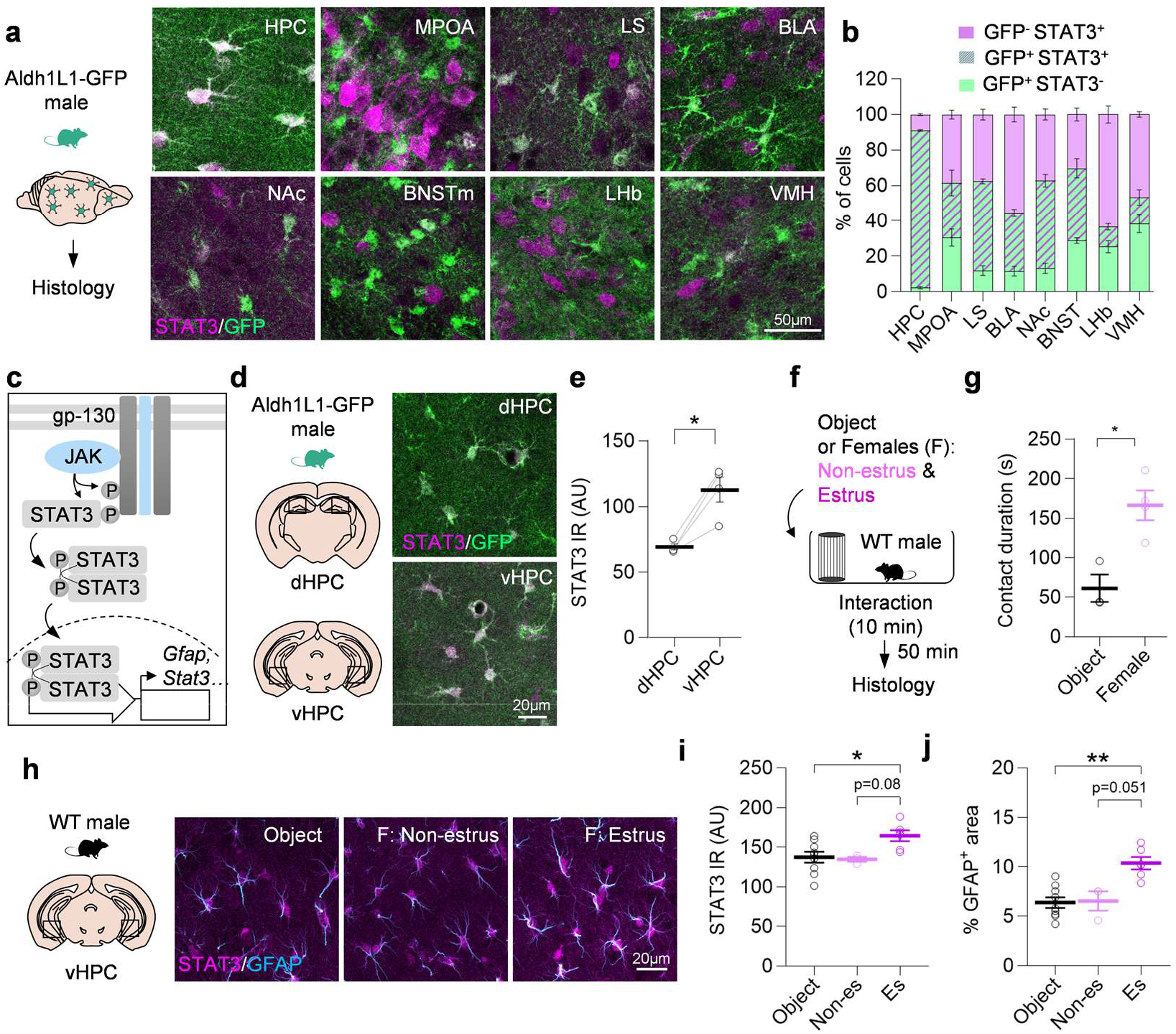
Exposure to receptive female induces astrocytic STAT3 signaling in the vHPC of male mice. **a,** Confocal images of brain sections from an astrocyte reporter mouse, Aldh1L1-GFP (green) co-stained with STAT3 (magenta) in limbic brain regions involved in social behavior. Note the co-localization between GFP and STAT3 in the HPC (cells appear white). Abbreviations: BLA: basolateral amygdala, BNST: bed nucleus of the stria terminalis, HPC: hippocampus, LHb: lateral habenula, LS: lateral septum, MPOA: medial preoptic area, NAc: nucleus accumbens, VMH: ventromedial hypothalamus. **b**, Quantification of the proportion of GFP-STAT3+, GFP+STAT3+ and GFP+STAT3-cells in each brain region (n=3-4 mice). **c**, Schematic representation of cytokine-associated canonical JAK-STAT3 signaling in astrocytes. Ligands bind to cytokine receptors containing the gp-130 subunit, triggering the transactivation of Janus Kinase (JAK), which in turn phosphorylates the receptor leading to the recruitment and phosphorylation of Signal Transducers and Activators of Transcription 3 (STAT3) proteins. Activated STAT3 proteins dimerize and translocate to the nucleus where they activate the transcription of target genes, such as *Gfap* and *Stat3* in astrocytes. **d**, High magnification images of Aldh1L1-GFP+STAT3+ astrocytes in the dorsal (d) and ventral (v) HPC. **e**, Quantification of nucleo-somatic STAT3 immunoreactivity (IR) in individual GFP+ astrocytes (n=4 mice). **f**, Adult WT male mice were let to approach with an object (Obj) or non-estrus (Non-es) or estrus (Es) females through a barred cage. After 50 min, the brains of male mice were processed for histology. **g,** Contact duration of male mice exposed to an object and a female (n=3-4 mice). **h**, Confocal images of vHPC brain sections co-stained for STAT3 (magenta) and GFAP (cyan). **i, j,** Quantification of nucleo-somatic STAT3 IR in GFAP+ astrocytes (**i**) and the percentage of GFAP+ image area (**j**) (n=3-9 mice/group). **b, i**: One-way ANOVA with Sidak’s multiple comparisons tests; **j**: Kruskal-Wallis test and Dunn’s multiple comparisons test; **e, g**: Two-tailed paired t-test. *, p<0.05, **, p<0.01. Values were plotted as mean ±SEM.

Cytokine-associated STAT3 signaling relies on ligand binding to multimeric receptor complexes (involving the gp-130 receptor), triggering the transactivation of Janus Kinase (JAK), which in turn phosphorylates the receptor leading to the recruitment and phosphorylation of STAT3 ^11^. Activated STAT3 proteins dimerize and translocate to the nucleus where they activate the transcription of target genes, such as *Gfap* and *Stat3* in astrocytes ^15,16^ (**Figure 1c**). To assess basal JAK-STAT3 pathway activity in astrocytes, we quantified STAT3 immunoreactivity in astrocyte nucleo-somatic region, as a proxy for activation since phospho-STAT3 cannot be reliably detected in astrocytes of the healthy mouse brain ^15,16^ (**Figure 1d**). Interestingly, while highly enriched in astrocytes in both the dorsal (d) and the ventral (v) HPC, STAT3 levels were significantly higher in the vHPC than in the dHPC, suggesting that a more active signaling at baseline in the ventral subregion (**Figure 1e**).

As opposed to the dHPC, which plays crucial roles in spatial learning and memory, the vHPC is comparatively more involved in the processing of emotional information elicited by stress but also social cues ^24^. To evaluate whether astrocytic STAT3 signaling could be induced after social stimulation, we exposed male mice to females, a stimulus that was previously shown to activate vHPC neurons in the male rodent brain ^25,26^ (**Figure 1f**). To uphold an ethologically-relevant setting, free-cycling females were used as stimuli and exposed to males through a barred cage to avoid copulation. We first conducted a time course experiment to determine the optimal interval between behavioral stimulation and euthanasia for histological analysis. Adult WT male mice were perfused at 30, 75 and 120 min after the end of a 10-min interaction with a female and their brains processed for histological detection of STAT3 and GFAP, to assess signaling activation (**Extended Data Fig.1d**). Both STAT3 and GFAP protein levels were transiently increased at 75 min post-social interaction before decreasing (**Extended Data Fig.1e-g**). We decided to use a 50 min interval corresponding to the ascending phase of the time course for subsequent analyses. Following this time course experiment, additional adult WT male mice were let to interact with either an object or a female of known estrus cycle phase (**Figure 1f**) and expectedly spent more time in contact with the barred cage containing a female than an object (**Figure 1g**). We observed that both STAT3 and GFAP levels were higher in the vHPC of male mice after interaction with females in estrus than in non-estrus and an object (estrus vs object: +20% for STAT3 and +62% for GFAP; estrus vs non-estrus: +22% for STAT3 and +58% for GFAP) (**Figure 1h-j**). These results were confirmed in a separate mouse cohort by western blotting (**Extended Data Fig. 1h-k**). As additional controls, we compared STAT3 and GFAP immunoreactivity between mice that interacted with an object or mice that stayed in their home cage (**Extended Data Fig. 1l**). There was no significant difference of STAT3 signaling activation in vHPC astrocytes after exposure to an object as compared to the home cage controls (**Extended Data Fig.1m-o**), suggesting that mouse handling and exploration of an object in a novel environment are not confounding factors explaining STAT3 activation after female exposure. To assess if STAT3 activation following female exposure was specific to astrocytes, we quantified the percentage of STAT3+GFAP+ cells in male mice after exposure to a female or an object. After female exposure, STAT3 was still broadly astrocyte-specific, since it is expressed in >90% of GFAP+ cells, suggesting that STAT3 signaling is only induced in this cell type following behavioral stimulation (**Extended Data Fig. 1p**).

Together, these results show that STAT3 signaling basal activation is prominent in vHPC astrocytes of male mice and further induced following exposure to a female, in an estrus cycle-dependent manner.

### Experimental activation of JAK2-STAT3 signaling induces estrus female-like approach towards non-receptive females

To determine whether STAT3 signaling activation after exposure to estrus female was correlative or causative of behavioral changes in male mice, we experimentally activated the pathway in vHPC astrocytes, using a viral approach. We first validated that following targeted intracerebral injections, astrocyte-specific adeno-associated viral vectors carrying a GFA-ABC1D (GFA) promoter and encoding GFP (AAV-GFP) indeed preferentially transduced the vHPC whereas a very limited number of GFP+ cells were detected in the dHPC (**Extended Data Fig.2a-c**). We next evaluated AAV-GFP transduction efficiency (83,9%) and specificity (99,5%) using co-immunostainings with cell type-specific markers (**Extended Data Fig.2d, e**). We also confirmed that the injection itself did not trigger GFAP induction by comparing the % GFAP+ area on vHPC brain sections from non-injected and AAV-GFP-injected mice (**Extended Data Fig.2f, g**). Finally, since AAV-encoded transgenes used in this study cannot be detected due to a lack of reliable antibodies, we systematically co-transduced astrocytes with AAVs encoding the transgenes of interest and a fluorescent protein. To confirm that co-injection of identical AAVs co-transduce the same astrocytes, we injected WT mice with a mix of AAVs encoding GFP and td-Tomato and processed samples for histology and FACS-based analysis (**Extended Data Fig.2h**). The majority of astrocytes indeed expressed both transgenes, as seen on confocal images (**Extended Data Fig.2i**) and quantified by cytometry (**Extended Data Fig.2j**).

Having validated our viral approach, we next determined whether experimentally activating STAT3 signaling in vHPC astrocytes would influence male approach behavior towards females. We took advantage of a previously validated genetic strategy to activate STAT3, relying on the expression of a constitutively active mutant form of the upstream kinase, JAK2 (JAK2ca)^16,17^. Adult WT males were injected in the vHPC with astrocyte-targeted AAV and encoding either *Jak2ca* + *Gfp* or *Gfp* only (**Figure 2a**). STAT3 nucleo-somatic levels were higher in astrocytes of mice injected with AAV-JAK2ca than in AAV-GFP controls, validating an efficient activation of the pathway (**Figure 2b, c**). Interestingly, this activation was of similar amplitude (+22%) of that observed after male exposure to estrus female (**Figure 1h, i**), suggesting that AAV-JAK2ca induces a moderate and biologically-relevant activation of the pathway.

**Figure 2.**
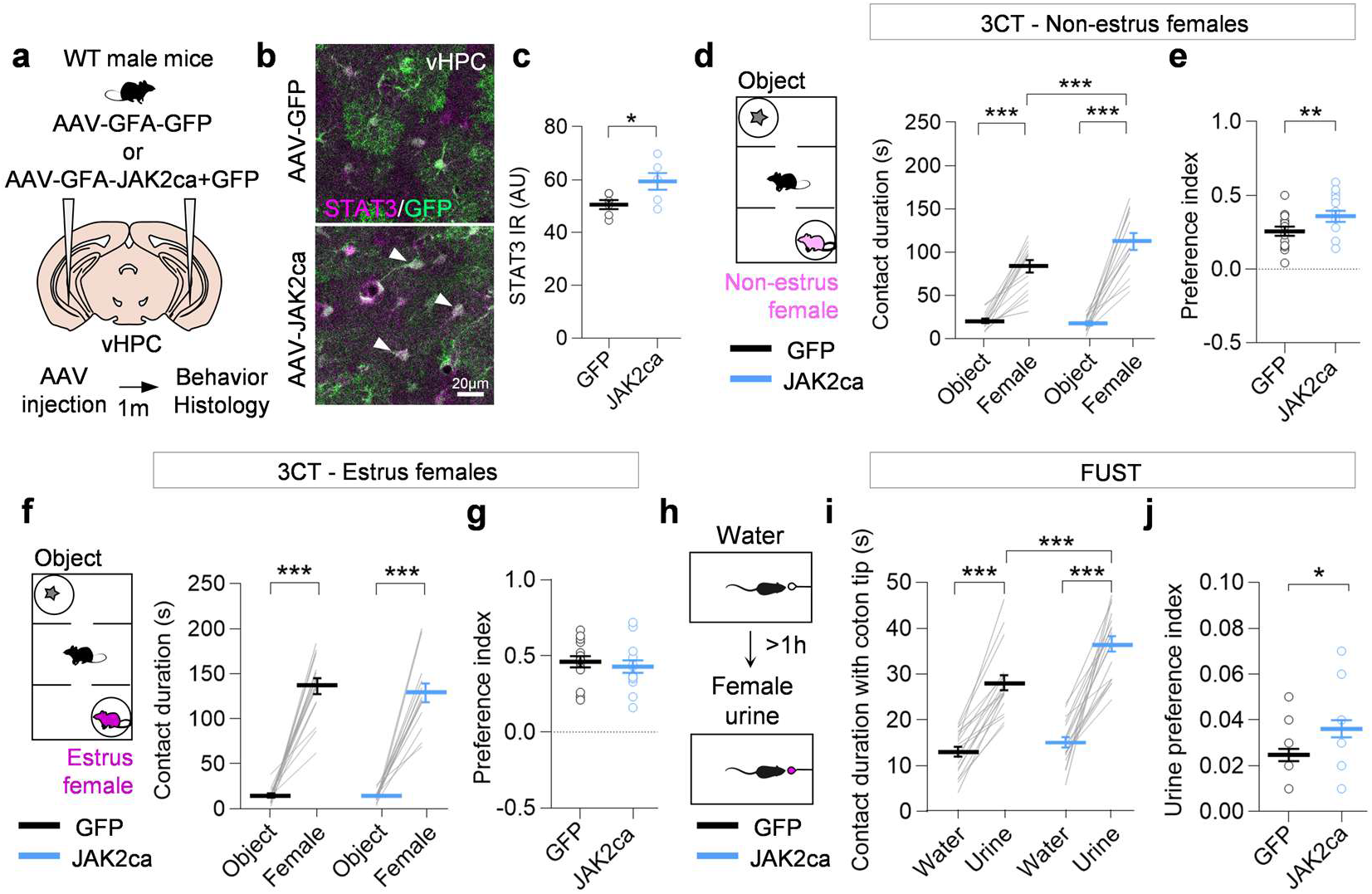
Experimental activation of vHPC astrocyte JAK2-STAT3 signaling induces estrus female-like male approach towards non-receptive females. **a,** Adult WT male mice received intracerebral injections of astrocyte-targeted AAV encoding *Jak2ca* or *Gfp* in the vHPC. **b,** High magnification confocal images of GFP (green)/STAT3 (magenta) co-immunofluorescent staining of brain sections from mice injected with AAV-JAK2ca and GFP controls. STAT3 nucleo-somatic localization is highlighted by white arrowheads. **c**, Quantification of nucleo-somatic STAT3 immunoreactivity in GFP+ astrocyte soma (n=6 mice/group). **d-g**, Sociability was assessed at the 3-chamber test with non-estrus (**d**) or estrus females (**f**), through the evaluation of contact durations (**d**, **f**) and preference indexes (**e**, **g**) (n=14-15 mice/group). **h**, AAV-JAK2ca and -GFP-injected mice were tested at the female urine sniffing test. **i, j,** Contact duration with cotton tip (**i**) and urine preference index (**j**) were evaluated (n=18-19 mice/group). **d**, **f**, **i**: Two-way repeated measure ANOVA and Sidak’s multiple comparison tests; **c**, **e**, **g**: Two-tailed unpaired t-test; **j**: Mann-Whitney test. *, p<0.05, **, p<0.01, ***, p<0.001. Values were plotted as mean ±SEM.

We then used the 3-chamber test to assess the effect of AAV-JAK2ca on male approach behavior towards females as compared to an object. When exposed to non-estrus females, AAV-JAK2ca males show increased contact duration and preference index for the female, as compared to GFP controls (**Figure 2d, e**). However, in response to estrus females, AAV-JAK2ca and -GFP mice display similar contact durations and preference indexes throughout the test (**Figure 2f, g**). To evaluate whether JAK2ca effect was specific to male-female encounters, we performed the test with juvenile mice (**Extended Data Fig.3a**). While AAV-JAK2ca and -GFP mice displayed a strong preference for the juvenile over the object, no group difference was observed (**Extended Data Fig.3b, c**). Furthermore, because of the well-characterized role of the vHPC in social memory ^24^, we also evaluated social novelty preference towards juveniles. Again, both AAV-JAK2ca and -GFP mice spent more time interacting with the novel versus familiar juvenile and no group difference was observed (**Extended Data Fig.3d, e**).

To determine if olfactory cues alone were sufficient to observe the enhanced preference for non-estrus females with AAV-JAK2ca mice, we performed the female urine-sniffing test. To do so, AAV-JAK2ca and -GFP-injected adult males were exposed to a cotton tip with water and at least 1h later, a cotton tip with female urine, as previously described ^27^ (**Figure 2h**). We found that the time in contact with female urine was increased in AAV-JAK2ca as compared to AAV-GFP mice, while not different for water (**Figure 2i, j**). We next assessed if the observed difference could arise from changes in olfactory processing and performed the olfactory habituation/dishabituation test with both social and non-social odors (**Extended Data Fig.3f, h**). We observed the correct pattern of olfactory habituation/dishabituation with each change of odor but there was no group effect between AAV-JAK2ca and -GFP mice, suggesting that they process olfactory cues similarly (**Extended Data Fig.3g, i**). Finally, we also assessed whether the experimental activation of the JAK2-STAT3 signaling in astrocytes also affected anxiety-like behavior, a well-described vHPC-related behavior ^24^. We did not find any significant difference in the performances of AAV-JAK2ca and -GFP mice at the elevated plus maze (**Extended Data Fig.3j-l**) and the open field test (**Extended Data Fig.3m-o**).

Together, these results suggest that activating the JAK2-STAT3 signaling in astrocytes differentially impacts approach behavior of male mice towards females depending on their receptivity.

### Astrocyte JAK2-STAT3 signaling influences vCA1 pyramidal neuron activity in response to female exposure

To understand the cellular circuit engaged downstream of astrocyte JAK2-STAT3 signaling, we investigated the impact of JAK2ca on neighboring pyramidal neurons. To do so, we first performed electrophysiological recordings on acute vHPC slices within the GFP+ area of AAV-JAK2ca+GFP or -GFP-injected mice (**Figure 3a, b, f**). Using whole-cell patch-clamp, we observed that vCA1 pyramidal neurons exhibited heightened basal excitability in the AAV-JAK2ca group as compared to AAV-GFP controls (**Figure 3c, d**) without alteration in input/output curves representing suprathreshold firing gain (**Figure 3e**). In line with this, evoked dendritic field excitatory postsynaptic potentials (fEPSPs) recorded in the vCA1 stratum radiatum and somatic population spikes recorded in the pyramidal cell layer following Schaffer’s collateral stimulation did not differ between groups (**Extended Data Fig.4a-d**). We further examined the effect of astrocyte JAK2-STAT3 activation on NMDAR-mediated responses. The input-output curve of NMDAR-fEPSPs in the AAV-JAK2ca group was significantly rightward shifted compared to AAV-GFP controls (**Figure 3g-i**). Since the pair-pulse ratio was comparable between groups (**Extended Data Fig.4c**); the rightward-shifted NMDAR-fEPSP and elevated AMPA/NMDA ratio (**Figure 3h, i**) suggested reduced postsynaptic NMDAR availability during synaptic activation without changes in presynaptic release probability.

**Figure 3.**
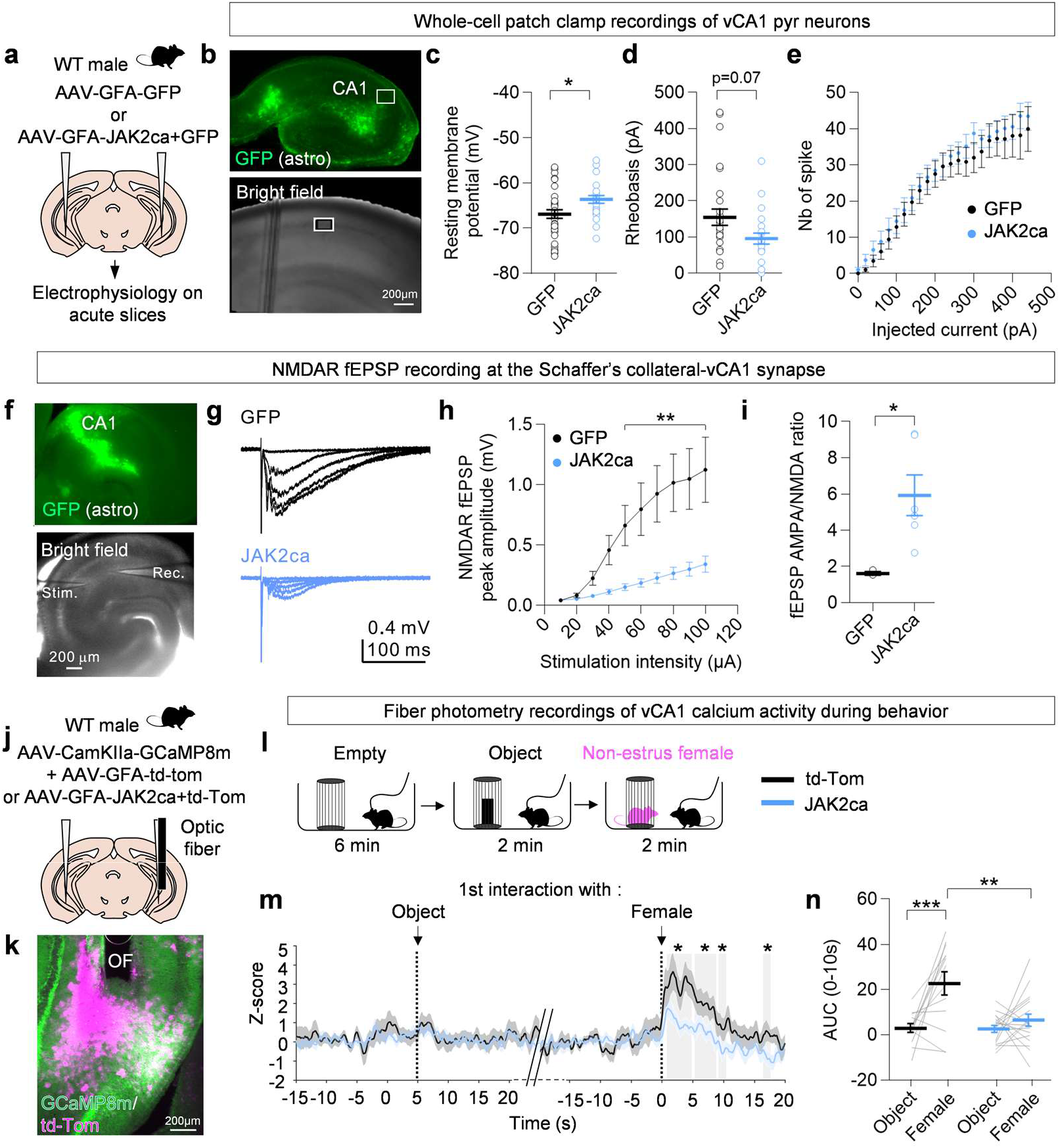
JAK2-STAT3 signaling activation in vHPC astrocytes influences neighboring vCA1 pyramidal neuron excitability and response to females. **a**, Adult WT male mice received intracerebral injections of astrocyte-targeted AAV encoding J*ak2ca* or G*fp* in the vHPC. Their brain was subsequently used for patch clamp recordings of vCA1 pyramidal neuron intrinsic properties on acute slices. **b**, Representative fluorescence (top) and brightfield (bottom) images of vHPC slices used for recordings. **c-e**, Quantification of vCA1 pyramidal neurons resting membrane potential (n=2-19 cells from 3 mice/group) (**c**), rheobasis (n=1-12 cells from 2-5 mice/group) (**d**) and input/output curves (n=2-18 cells from 3 mice/group) (**e**) comparing neurons from the JAK2ca and GFP control groups. **f**, NMDAR fEPSP recording at the Schaffer’s collateral-vCA1 synapse in the GFP+ area of AAV-injected mice. **g**, Example traces of NMDAR fEPSP recordings from GFP (black) and JAK2ca (blue) acute slices. **h, i**, NMDAR fEPSP peak amplitude (**h**) and AMPA/NMDA fEPSP ratio (**i**). **j**, Adult WT males received bilateral intracerebral injections of a mix of an AAV targeting neurons and encoding the genetic Ca2+ indicator *Gcamp8m* and of AAVs targeting astrocytes enabling the expression of J*ak2ca* and/or *td-Tomato*. Mice were implanted in the vHPC with an optic fiber to subsequently perform photometry recordings. **k**, Low magnification image of the vHPC of a mouse expressing GCaMP8m (green) and td-Tomato (magenta). The optic fiber tract is noted as (OF). **l**, Implanted male mice were exposed to an empty barred cage for 6 min (baseline), followed by exposure to an object for 2 min and a female for 2 min, during which Ca2+ activity of vCA1 neurons was recorded. **m**, Z-score plot showing the Ca2+ activity of vCA1 neurons after interaction with an object than with a female in both JAK2ca and td-Tomato control groups. Grey blocks indicate the time window with a significant signal difference between AAV-JAK2ca and -GFP groups using a permutation test (n=13-18 mice/group). **n**, Area under the curve (AUC) was calculated on Z-score plot showed in **m**. **c**, **i**: Two-tailed unpaired t-test; **d**: Mann-Whitney test; **e**: Mixed model with Geisser-Greenhouse’s correction and Sidak’s multiple comparison test; **h, n**: Two-way repeated measure ANOVA and Sidak’s multiple comparison test. **m**: Two-sided permutation test. * p < 0.05, **p < 0.01, ***p < 0.001. Values were plotted as mean ±SEM.

To determine whether our *ex vivo* observations would translate into changes in vCA1 pyramidal neurons response to a behavioral stimulation, we co-injected pyramidal neuron-targeted AAV encoding the genetically-encoded Ca2+ indicator *Gcamp8m* and astrocyte-targeted AAV encoding either *Jak2ca* + *td-Tomato* or *td-Tomato* in the vHPC of male mice. Animals were also implanted in the vHPC with an optic fiber cannula to record Ca2+ activity of pyramidal neurons during behavioral stimulation (**Figure 3j**). Cannula implantation along with GCaMP8m and td-Tomato transgenes expression were controlled using immunofluorescence (**Figure 3k**). Calcium activity was recorded in vCA1 pyramidal neuron of WT males sequentially exposed to an empty barred cage, as baseline, followed by the same cage containing an object and finally a non-estrus female (**Figure 3l**). The normalized GCaMP8m fluorescence signal was similar during interaction with the object as compared to baseline, in both AAV-JAK2ca and -td-Tomato control groups (**Figure 3m, n**). By contrast, we observed sharp signal increase after the first contact with the female as compared to baseline, in both groups although this response was lower in the AAV-JAK2ca than in the AAV-td-Tomato group (**Figure 3m, n**). At the end of each recording session, tail suspension was performed for 10 sec. Given that this stressful stimulation led to a prominent increase in vCA1 Ca2+ signals although with the same amplitude in both the AAV-JAK2ca and -td-Tomato controls (**Extended Data Fig.4f-h**), we used it as a positive control for signal quality assessment.

Altogether, these findings suggest that continuous astrocytic JAK2-STAT3 activation drives basal hyperexcitability of vHPC neuronal network while impairing NMDAR-mediated synaptic transmission, thereby blunting female-evoked neuronal Ca2+ dynamics during social behavior.

### Chemogenetic activation of vCA1 pyramidal neurons enhances male approach towards non-estrus females

We then asked whether directly activating vCA1 neurons could replicate JAK2ca-induced behavioral changes. To do so, we used a chemogenetic approach by injecting WT male in the vHPC with an activating DREADD (hM3Dq-Cherry) targeting pyramidal neurons (CaMKII promoter). One-month post-intracerebral injections, mice were administered with Clozapine-N-oxide (CNO) (0.1 mg/kg) or vehicle (VEH) and underwent histological analysis and behavioral testing 1h post-administration (**Figure 4a**). By quantifying the percentage of c-Fos+mCherry+/total mCherry+ cells, we confirmed that CNO administration induced c-Fos expression- a proxy for cellular activation-in ∼90% of transduced vCA1 pyramidal neurons (**Figure 4b, c**).

**Figure 4.**
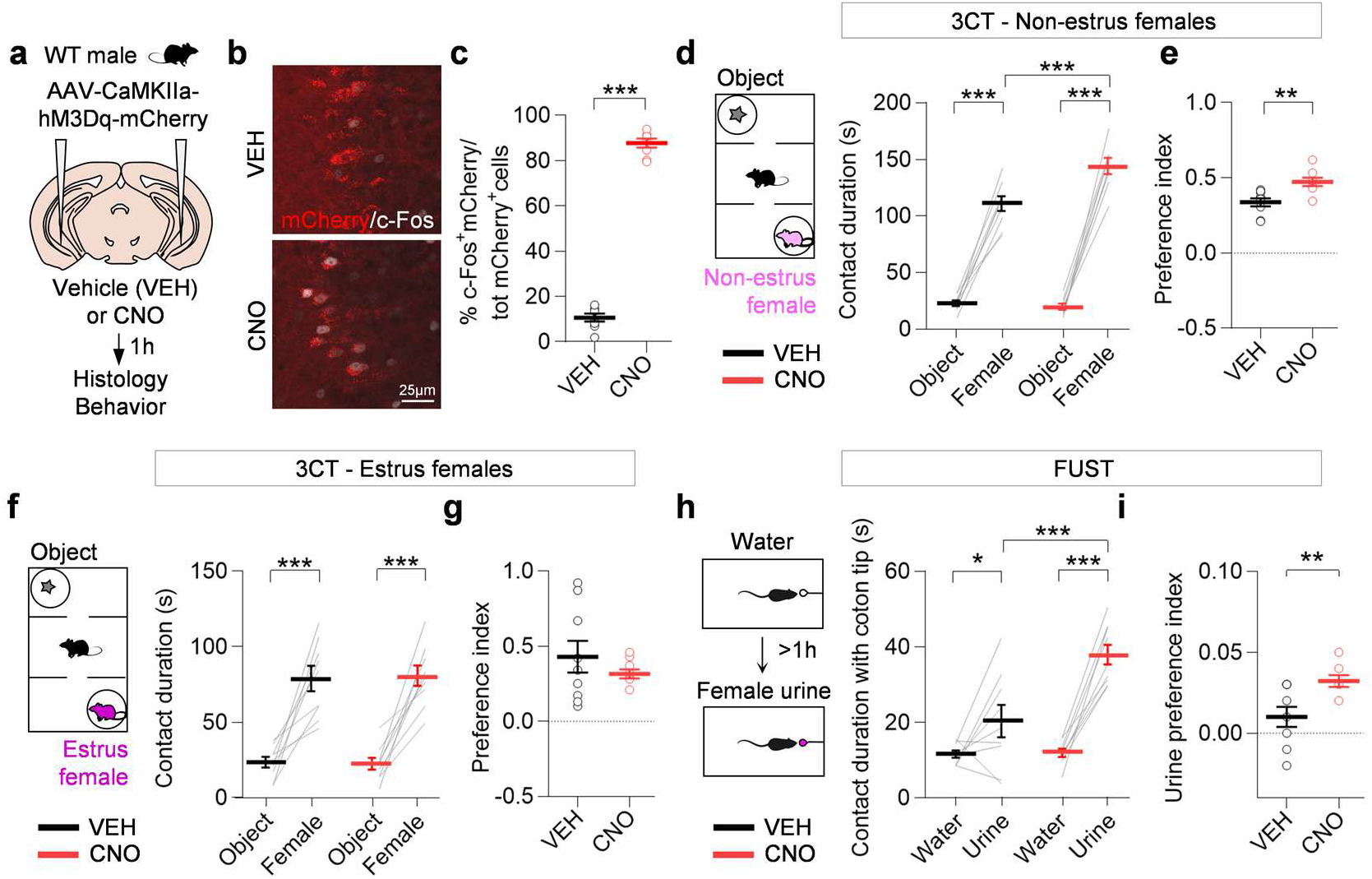
Chemogenetic activation of vCA1 pyramidal neurons enhances male approach towards non-estrus females. **a,** Adult WT male mice were injected in the vHPC with AAVs targeting pyramidal neurons (CamKIIa promoter) and encoding *hM3Dq-mCherry*. One-month post-injection, mice were administered with CNO (0.1 mg/kg) or vehicle (VEH) via micropipette-assisted drug administration one hour prior to histological analysis or behavioral testing. **b**, Confocal images of vHPC brain sections from AAV-CamKIIa-hM3Dq-mCherry-injected mice after administration of VEH or CNO stained for c-Fos (gray) and mCherry (red). **c**, Quantification of the % of c-Fos+mCherry+/total mCherry+ cells (n=8 mice/group). **d**-**g**, Sociability was assessed at the 3-chamber test with non-estrus females (**d**) or estrus females (**f**), through the evaluation of contact durations (**d**, **f**) and preference indexes (**e**, **g**) (n=8-9 mice/group). **h-i**, Contact duration with water and female urine (**h**) and preference index (**i**) (n=9 mice/group). **c**, **e**, **g**, **i**: Two-tailed unpaired t-test, **d**, **f**, **h**: Two-way repeated measure ANOVA and Sidak’s multiple comparison tests. *p < 0.05, **p < 0.01, ***p < 0.001. Values were plotted as mean ±SEM.

We next investigated AAV-CaMKIIa-hM3Dq-injected male approach behavior at the 3-chamber test with non-estrus (**Figure 4d**) and estrus (**Figure 4f**) females against an object, 1h after CNO or VEH administration. Mice from the CNO group displayed significantly higher contacts with non-estrus but not with estrus females, while their exploration of the object was similar to the VEH controls (**Figure 4d-g**). Moreover, at the female urine sniffing test, CNO-administered mice also showed an increased contact duration as compared to mice in the VEH group (**Figure 4h, i**). Since CNO may cause off-target effects by acting on endogenous receptors^39^, we performed control experiments to validate that the molecule itself did not impact the performances of male mice at the 3-chamber test. To do so, we repeated the last experiments in AAV-CaMKIIa-GFP control mice (**Extended Data Fig.5a**). At the histological level, the number of c-Fos+ cells was similar between CNO and VEH groups (**Extended Data Fig.5b, c**). At the behavioral level, we did not observe significant group differences at the 3-chamber test with non-estrus females (**Extended Data Fig.5d, e**) and at the female urine sniffing test (**Extended Data Fig.5f, g**).

Together, these results suggest that the direct activation of vCA1 pyramidal neurons enhances male approach behavior towards non-estrus females, thus replicating the effect observed by expressing JAK2ca in neighboring vHPC astrocytes.

### Inhibiting cytokine-associated JAK2-STAT3 signaling in vHPC astrocytes does not influence male-to-female approach behavior

We then asked whether blocking astrocytic JAK2-STAT3 signaling in the vHPC would mirror the behavioral effects observed with JAK2ca i.e., decreased male approach towards non-receptive females. We used a viral approach to overexpressed SOCS3, the endogenous inhibitor of the JAK2-STAT3 pathway ^28^(**Figure 5a**). WT male mice received intracerebral injections of astrocyte-targeted AAV encoding either *Socs3* and *Gfp* or *Gfp* only (**Figure 5b**) as previously described ^15–17^. STAT3 immunoreactivity was significantly reduced after injection of AAV-SOCS3 as compared to AAV-GFP controls, confirming that overexpression of SOCS3 successfully inhibits downstream JAK2-STAT3 signaling (**Figure 5c, d**). We next evaluated approach behavior AAV-SOCS3 and -GFP mice at the 3-chamber test. Interestingly, we did not observe any significant differences in the contact duration between groups with both non-estrus and estrus females (**Figure 5e-h**), suggesting that the canonical JAK2-STAT3 signaling is not required for normal male-to-female approach behavior.

**Figure 5.**
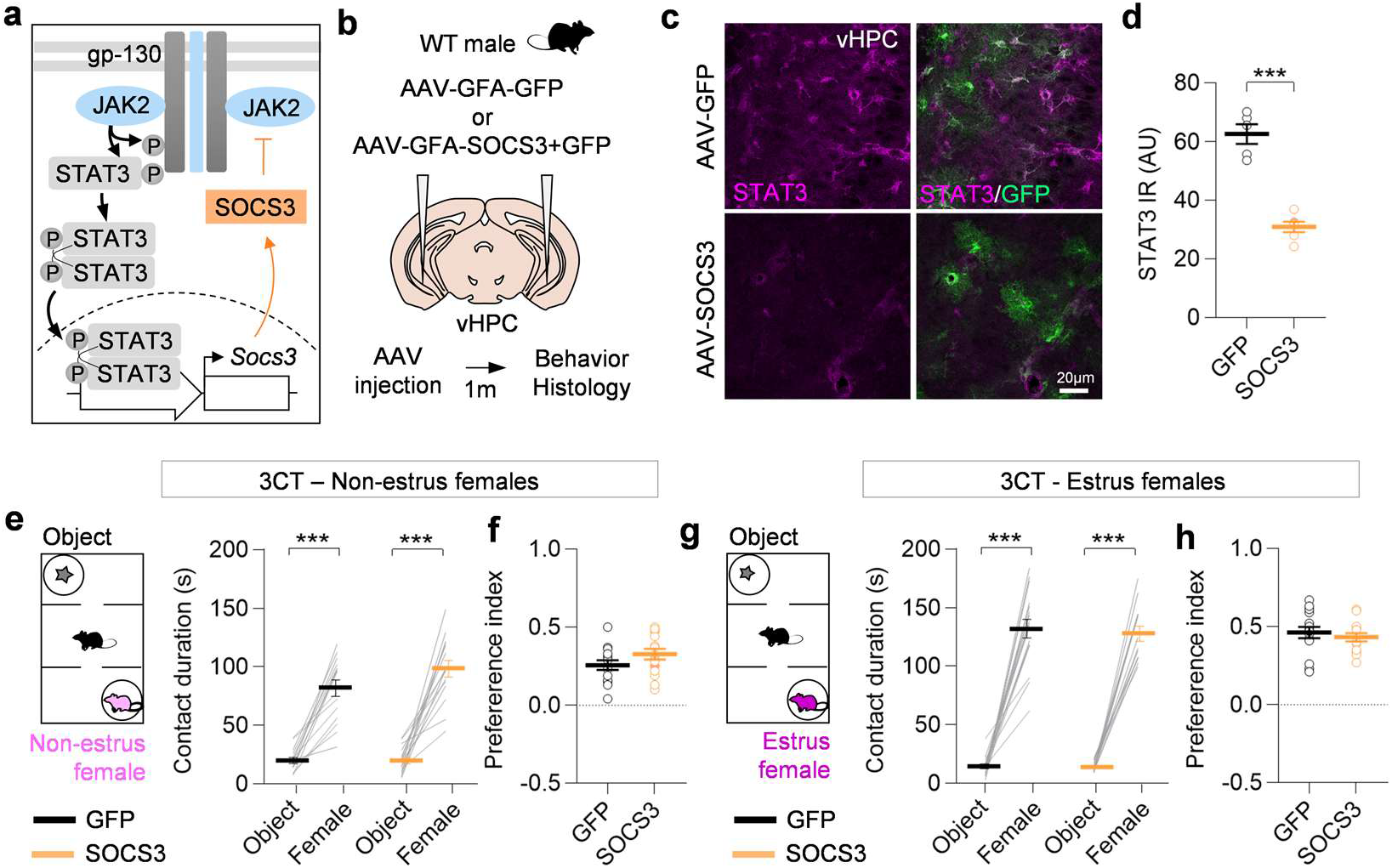
Inhibition of cytokine-mediated JAK2-STAT3 signaling in vHPC astrocytes does not influence male-to-female approach behavior. **a,** Schematic representation of cytokine-associated JAK2-STAT3 signaling in astrocytes. The JAK2-STAT3 signaling is regulated through an inhibitory feedback loop, via the induction of SOCS3, which blocks JAK2 phosphorylation. **b,** Adult WT male mice were injected with AAV targeting astrocytes and encoding *Gfp* as control or *Socs3*+*Gfp* to block the JAK-STAT3 signaling. One-month post-injection, mice underwent behavioral testing and were subsequently euthanized to analyze their brain by histology. **c**, Confocal images of vHPC brain sections co-immunofluorescently labeled with STAT3 (magenta) and AAV-encoded GFP (green). **d**, Nucleo-somatic STAT3 immunoreactivity (IR) quantification in AAV-GFP and -SOCS3 groups (n=5-6 mice/group). **e-h**, AAV-injected mice were tested at the 3-chamber test with non-estrus (**e, f**) and estrus (**g, h**) females against an object (n=14-15 mice/group). The contact duration with each stimulus and the preference index are shown. **d, f**, **h**: Two-tailed unpaired t-test; **e**, **g**: Two-way repeated measure ANOVA with Sidak’s multiple comparison test. ***, p < 0.001. Values were plotted as mean ±SEM.

Given that SOCS3 blocks the signaling through binding of JAK2 when recruited to the gp130 receptor^29^, we hypothesized that a parallel, gp130 receptor-independent, upstream activator may induce astrocytic JAK2-STAT3 signaling upon receptive female exposure.

### Activation of septo-hippocampal cholinergic neurons upon estrus female exposure recruits astrocytic JAK-STAT3 signaling in the vHPC

Previous work showed that the vHPC receives cholinergic innervation from the medial septum (MS) and the Diagonal Band of Broca (DBB) ^30,31^. Given that acetylcholine (ACh) anti-inflammatory roles have been linked to the JAK2-STAT3 signaling in peripheral immune cells ^32^, we surmised that vHPC astrocytic JAK2-STAT3 signaling could be activated downstream of ACh.

To confirm vHPC cholinergic innervation, we used a genetic labeling strategy by crossing ChAT-IRES-cre mice with td-Tomato Cre reporter mice (Ai9) leading to td-Tomato expression in choline acetyltransferase (ChAT)+ cells, that we validated using co-immunofluorescent stainings (**Figure 6a-c**). We indeed observed td-Tomato+ projections in the vHPC and cell bodies in the MS/DBB, in accordance with previous findings (**Figure 6b, c**). To further confirm the MS/DBB-to-vHPC cholinergic innervation, adult ChAT-IRES-Cre mice were injected in the vHPC with a Cre-dependent retro-AAV encoding mCherry (**Extended Data Fig.6a**). After local injection of retro-AAV in the vHPC, we observed mCherry+ axon terminals in the vHPC and retrogradely labelled ChAT+ cell bodies in the MS/DBB (**Extended Data Fig.6b-d**). These experiments validated that the vHPC receives cholinergic inputs from the MS/DBB in the adult male mouse brain.

**Figure 6.**
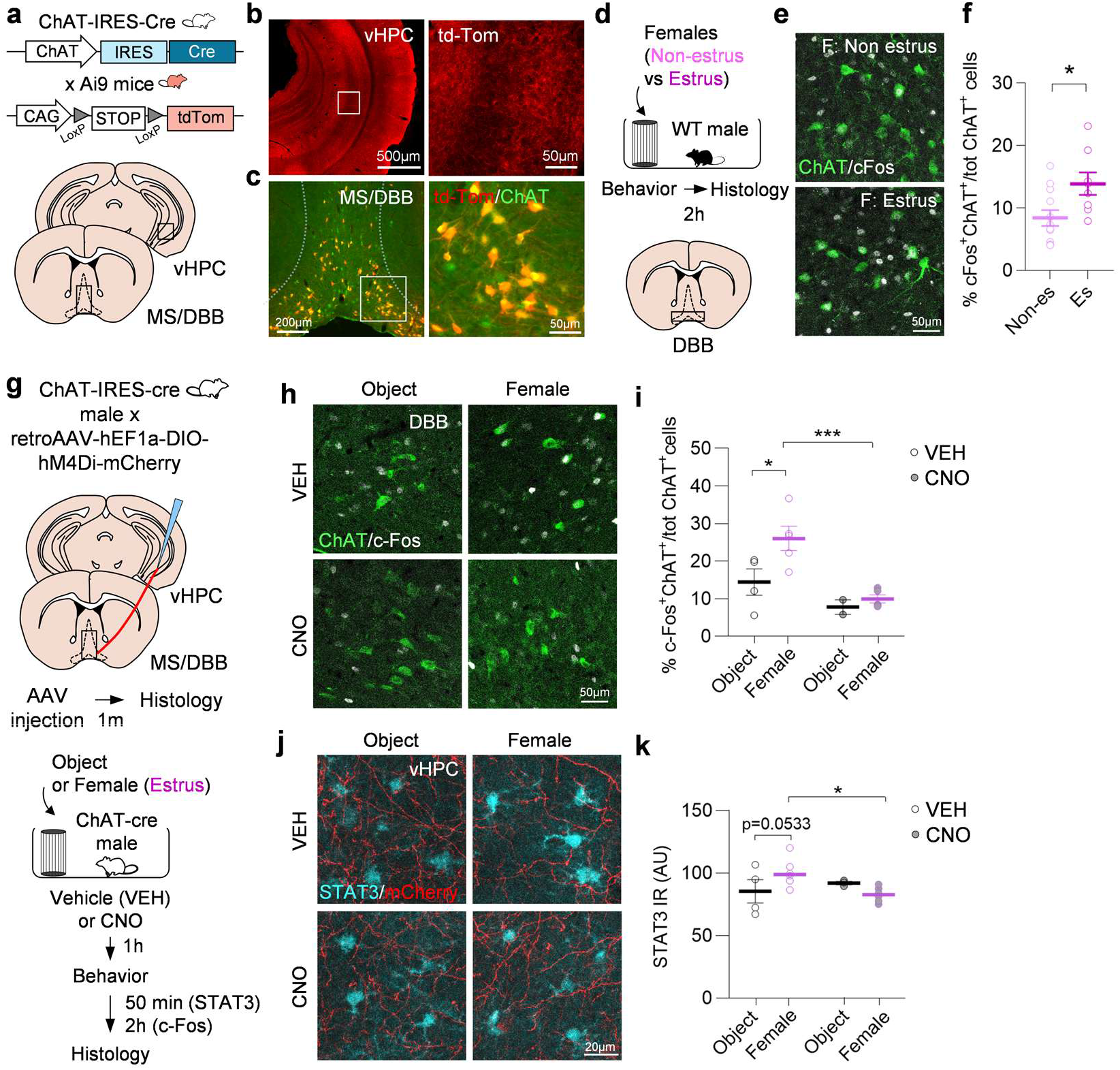
Septo-hippocampal cholinergic neuron activation induces the JAK2-STAT3 signaling in vHPC astrocytes. **a,** Ai9 td-Tomato reporter mice were bred with ChAT-IRES-Cre mice to target Cre-dependent expression of td-Tomato to ChAT+ cells. **b,** Images of td-Tomato labeling in the vHPC at low (right) and high (left) magnification. **c,** High magnification confocal images of MS/DBB brain sections co-stained for td-Tomato (red) and ChAT (green). **d**, WT male mice were let to interact with non-estrus and estrus females for 10 min and their brains were analyzed by histology 2h later. **e**, Confocal images of DBB brain sections after interaction with an non-estrus (top) and estrus (bottom) females, co-stained ChAT (green) and c-Fos (gray). **f**, Quantification of the % c-Fos+ChAT+/total ChAT+ cells in DBB after interaction with non-estrus and estrus females (n=8-12 mice/group). **g**, ChAT-cre male mice received intracerebral injections of a cre-dependent retrograde AAV in the vHPC encoding the inhibitory DREADD, hM4Di-mCherry. Mice were administered either 2 mg/kg of clozapine-N-oxide (CNO) or vehicle (VEH) and were behaviorally tested 1h later. Independent cohorts were then euthanized 50 min (for STAT3 histology) or 2h (for c-Fos histology) after behavior. **h**, Confocal images of DBB brain sections co-stained for ChAT (green) and c-Fos (gray). **i**, Quantification of the % c-Fos+ChAT+/total ChAT+ in the DBB after interaction with an object or a female, in the CNO and VEH groups (n=2-5 mice/group). **j**, Confocal images of vHPC brain sections co-stained for mCherry (red) and STAT3 (cyan). **k**, Quantification of nucleo-somatic STAT3 immunoreactivity (IR) after interaction with an object or a female, in the CNO and VEH groups (n=3-6 mice/group). **f**: Two-tailed unpaired t-test; **i**, **k**: Two-way ANOVA and Sidak’s multiple comparison tests. *, p < 0.05, ***, p < 0.001. Values were plotted as mean ±SEM.

We next determined whether male exposure to female recruited septal cholinergic neurons by performing co-immunostaining for ChAT and c-Fos in the MS and DBB of male mice after exposure to a female or an object (**Extended Data Fig.6e**). Interestingly, while the number of c-Fos+ cholinergic neurons was systematically higher in the DBB as compared to the MS, it was only significantly induced in the DBB after female exposure (**Extended Data Fig.6f, g**). We next compared the activation of DBB cholinergic neurons following male approach between estrus and non-estrus females (**Figure 6d**). Consistent with our previous findings, the percentage of ChAT+/cFos+ neurons in the DBB was higher following male exposure with estrus than non-estrus females (**Figure 6e, f**).

To determine whether the activation of MS/DBB cholinergic neurons is required for the induction astrocyte JAK-STAT3 signaling within the vHPC, we employed an intersectional chemogenetic approach to inhibit MS/DBB cholinergic inputs onto the vHPC. To do so, we express the inhibitory DREADD, hM4Di-mCherry, in vHPC-projecting ChAT+ neurons by injecting ChAT-IRES-Cre male mice in the vHPC with a Cre-dependent (DIO) retro-AAV (**Figure 6g**). One hour after CNO (2 mg/kg) or VEH administration, AAV-DIO-hM4Di-mCherry injected males were exposed to an estrus female or an object for 10 min. Consistent with our previous findings, while the proportion of c-Fos+ cholinergic neurons was higher after female exposure in the VEH control group, it was significantly reduced following CNO administration (**Figure 6h, i**). We then quantified STAT3 immunoreactivity in mCherry+ area on vHPC sections from ChAT-IRES-Cre male mice injected with AAV-retro-DIO-hM4Di-mCherry (**Figure 6j**). Although values for STAT3 upregulation were just below significance, its levels appear to be increases after interaction with a female as compared to an object in the VEH condition, in accordance with our previous findings (**Figure 6k**). By contrast, CNO administration significantly reduced STAT3 levels selectively after male encounter with a female but not an object (**Figure 6k**). To control that CNO itself did not trigger JAK-STAT3 signaling activation, we detected STAT3 and GFAP on vHPC brain sections from non-injected WT male mice after administration of CNO (2 mg/kg) and VEH (**Extended Data Fig.7a**). We determined that both proteins were expressed at the same level in VEH and CNO conditions, therefore ruling out a non-specific effect of the CNO on JAK2-STAT3 signaling in vHPC astrocytes (**Extended Data Fig.7b-d**).

Taken together, these results suggest that MS/DBB ChAT+ cholinergic neurons projecting to the vHPC are activated after exposure to estrus female, presumably leading to a local release of ACh, which activates astrocyte JAK2-STAT3 signaling.

### Astrocyte α7nAChR signals through the JAK2-STAT3 signaling and modulates adapted male-to-female approach behavior

Given that JAK2 has been shown to interact with the α7 subunit of nicotinic acetylcholine receptors (α7nAChR) in peripheral immune cells ^33,34^, we hypothesized that α7nAChR-JAK2-STAT3 signaling also occurs in astrocytes of the adult mouse brain (**Figure 7a**).

**Figure 7.**
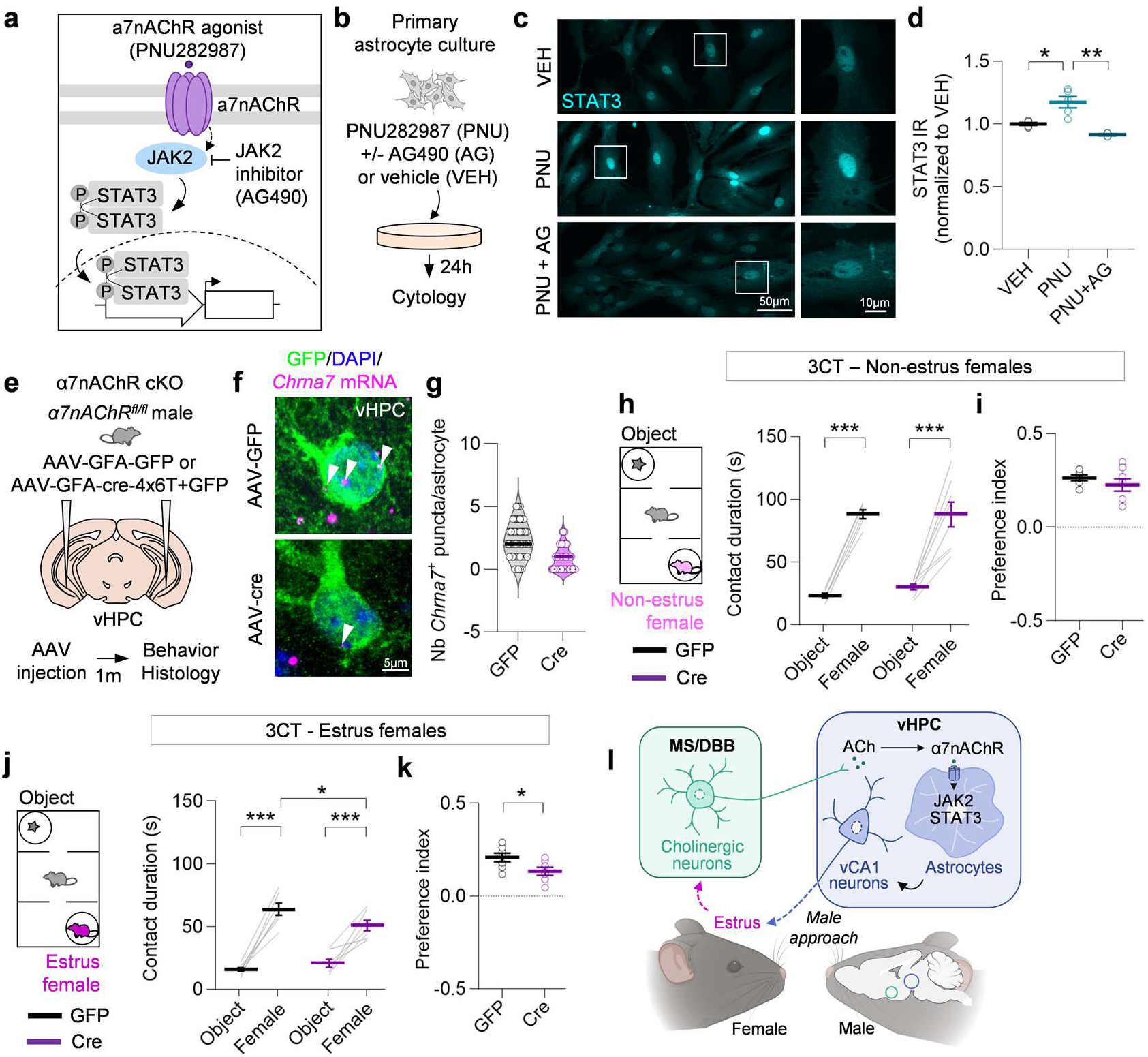
Astrocyte α7nAChR signals through the JAK2-STAT3 signaling and modulates adapted male-to-female approach behavior. **a,** Schematic of the cascade implicating JAK2-STAT3 signaling downstream of α7nAChR. Acetylcholine binds to α7nAChR, which activates JAK2, which in turn phosphorylates STAT3. Activated STAT3 proteins translocate to the nucleus where they induce the transcription of target genes. The α7nAChR-specific agonist PNU282987 and the JAK2 inhibitor AG490 are used to modulate this axis. **b,** Primary astrocyte cultures were treated 24h with 10µm PNU282987 only (PNU), in combination with 30µm AG490 (PNU+AG), or with the vehicle (VEH), DMSO for 24h and then fixed for immunocytology. **c,** Epifluorescence microscope images of STAT3 immunostaining (cyan). **d**, Normalized quantification of STAT3 nuclear signal (n=4-5 wells from 2 cultures). **e,** Adult male *α7nAChR^fl/fl^*mice were injected in the vHPC with AAV targeting astrocytes and encoding either a control *Gfp* or *Cre* + *Gfp*. One-month post-injection, mice were used for further processing. **f,** Co-detection of virally-encoded GFP protein (green) by immunofluorescence, α7nAChR-encoding mRNA (*Chrna7*) (magenta) by fluorescent *in situ* hybridization and DAPI on brain sections from *α7nAChR^fl/fl^* mice injected with AAV-GFP control (top) and -Cre (bottom). **g**, Quantification of the number of *Chrna7^+^* puncta per GFP^+^ astrocyte in the AAV-GFP and -Cre groups (n=50-51 cells from 3 mice). **h-k**, AAV-injected *α7nAChR^fl/fl^*mice underwent behavioral testing at the 3-chamber test with non-estrus (**h**) and estrus (**j**) females. Contact durations and preference indexes with an object versus the non-estrus (**h, i**) and estrus (**j, l**) females were measured (n=6-8 mice/group). **l**, Summary schematic. Estrus female exposure induces in the brain of male mice the recruitment of MS/DBB cholinergic inputs to activate vHPC astrocytic JAK2-STAT3 signaling downstream of α7nAChR, ultimately modulating adaptive approach behavior through changes in vCA1 pyramidal neuron activity. **d:** Linear mixed model analysis (fixed effect: treatment; random effect: well); **h**: Linear mixed model analysis (fixed effect: group; random effect: mouse); **h**, **j**: Two-way repeated measure ANOVA and Sidak’s multiple comparison tests; **i**, **k**: Two-tailed unpaired t-test. * p < 0.05, ** p < 0.01, ***p < 0.001. Values were plotted as mean ±SEM.

To test this hypothesis, we first used a pharmacological approach on pure murine primary astrocyte cultures, in which 100% of DAPI+ cells also expressed the astrocyte-specific markers phosphoglycerate dehydrogenase (PHGDH) and S100b (**Extended Data Fig.8a-c**). Primary astrocyte cultures were incubated for 24h with the α7nAChR agonist PNU282987, with or without a JAK2-specific inhibitor (AG490) or VEH followed by STAT3 immunodetection (**Figure 7b**). STAT3 nuclear levels were significantly increased in PNU282987-treated cells as compared to VEH, which was blocked by AG490 (**Figure 7c, d**). This result shows that STAT3 activation following α7nAChR pharmacological stimulation requires JAK2 in murine astrocytes, suggesting that these cells can signal through an α7nAChR-JAK2-STAT3 axis.

Based on these results, we then determined whether interfering with α7nAChR would mirror JAK2ca behavioral effects on male approach behavior towards females, depending on their receptivity. To do so, we used a cell type-specific conditional knockout (cKO) of α7nAChR-encoding gene (*Chrna7*) in astrocytes (**Figure 7e**). Mice were injected in the vHPC with an AAV targeting astrocytes and encoding a newly-developed Cre recombinase vector containing multiple detargeting sequences (4×6T) to repress potential ectopic Cre expression in neurons and GFP to visualize transduced astrocytes ^35^. Injection of AAV-Cre in Ai9 reporter mice led to a recombination efficiency of ∼100%, as assessed by the co-expression of GFP and td-Tomato in cells expressing the astrocyte marker, GS (**Extended Data Fig.9a-d**). In α7nAChR ^fl/fl^ mice, injection of AAV-Cre decreased the number of astrocyte *Chrna7+* mRNA puncta as compared controls, although it was not completely abolished (**Figure 7g, h**). By contrast, the number of *Chrna7+* puncta in neighboring pyramidal neurons was not different between AAV-Cre and -GFP controls (**Extended Data Fig.9e-g**). We then evaluated the effect of α7nAChR cKO in vHPC astrocytes on approach behavior of male mice towards females at the 3-chamber test. While AAV-Cre mice had a similar interaction with non-estrus females as compared AAV-GFP controls, they displayed a lower duration of contact and preference index for estrus females (**Figure 7h-k**), suggesting that the astrocytic downregulation of α7nAChR in the vHPC impairs normal behavior of male mice towards females, in an estrus cycle phase-dependent manner.

## Discussion

Our study reveals a physiological role for the JAK2-STAT3 pathway-a canonical mediator of astrocyte reactivity and neuroinflammation^36^- in astrocyte-neuron communication. We demonstrate that male exposure to estrus females rapidly and selectively induces STAT3 signaling in vHPC astrocytes. This induction depends on septo-ventral hippocampal cholinergic inputs acting through α7nAChRs, and in turn modulates vCA1 pyramidal neuron activity to shape adaptive male approach behavior towards females (**Figure 7l**). These findings establish an astrocyte-specific pathway that integrates neuromodulatory cues with TF-based signaling to gate male behavioral responses according to female reproductive state.

Transcription factors contribute to astrocyte regional specialization through the establishment of distinct molecular profiles ^37,38^. More recent work also uncovered that TFs such as SOX9 and c-Fos can be dynamically induced in an activity-dependent manner to regulate neuronal circuits involved in olfactory processing and memory recall, with evidence linking neuromodulation (via noradrenaline and serotonin) to TF-based signaling ^9,10,39,40^. Our results reinforce an emerging framework in which neuromodulatory gating of transcriptional pathways might represent a general signal transduction principle in astrocytes, triggered by behavioral stimulation. Such mechanisms likely prime astrocyte-neuron crosstalk to subsequent stimuli, thereby supporting behavioral adaptation. Notably, social cues probably engage multiple parallel astrocytic cascades, including Ca2+ signaling ^41–43^. Whether the JAK2-STAT3 signaling intersects with these alternative pathways remains an open question.

Our work also advances understanding of vHPC cholinergic modulation in male socio-sexual behavior by revealing a critical astrocytic contribution. Although a causal role for ACh in male sexual behavior was only recently described in the nucleus accumbens ^44^, we show that blocking astrocytic α7nAChR signaling in the vHPC reduces male contact time with receptive females. While we cannot generalize the effects of astrocytic manipulation on sexual behavior, our observations point to changes in male approach, an essential precursor to copulation. Moreover, because α7nAChR downregulation likely also impacts α7nAChR-mediated Ca²⁺ signaling in vHPC astrocytes, the observed behavioral phenotype likely reflects combined effects on multiple concurrent astrocytic pathways, warranting further investigation ^41^. Importantly however, our experiments with constitutively active JAK2 selectively alter male behavior toward females but not towards juveniles and does not impact other vHPC-dependent behaviors, supporting a specific role for the α7nAChR-JAK2-STAT3 axis in socio-sexual responses.

This study demonstrates a causal role for vHPC astrocytes in controlling male approach behavior towards females by modulating vCA1 pyramidal neuron activity. Previous work showed that vCA1 neurons selectively respond to social encounters and olfactory cues ^25,26^. Here, we find that activating astrocytic JAK2-STAT3 signaling during male-to-female interaction dampens vCA1 neuronal responses using in vivo fiber photometry. Although we cannot rule out that baseline signal could be higher in the JAK2ca group, this effect was not observed during tail suspension, suggesting that it is specific to social interaction. Additionally, our *ex vivo* fEPSP recordings, showing that JAK2-STAT3 activation does not influence basal synaptic transmission but decreases NDMAR-mediated response to stimulation, which might be a compensatory mechanism to prevent circuit overexcitation during behavioral stimulation in the JAK2ca group. In parallel, patch-clamp recordings suggest that JAK2-STAT3 activation in astrocytes enhanced vCA1 pyramidal neuron excitability. We thus hypothesize that female exposure elicits a reduced response in otherwise hyperexcitable vCA1 pyramidal neurons-an effect sufficient to facilitate male approach.

Collectively, our data uncover a new role for α7nAChR-JAK2-STAT3 signaling in astrocytes in the physiological regulation of neural circuits. These findings support a growing view that TF-based cascades are not merely static scaffolds for astrocyte molecular diversity or regulators of reactive changes, but dynamic signal transduction pathways central to the astrocyte-neuron crosstalk. Furthermore, α7nAChR-JAK2-STAT3 signaling may be co-opted by neuroinflammatory signals in disease, preventing astrocyte-mediated control of brain function in physiology.

## Acknowledgements

We thank all the Escartin lab members for technical help, feedback and advice. We thank MIRCen’s (Université Paris-Saclay, CEA, CNRS) viral core (A. Bemelmans, D. Fourmy), animal experimentation (G. Auregan, V. Letenneur), behavioral (K. Cambon) and histology (F. Petit, P. Giptchein) platforms. We thank NeuroPSI’s NeuroPICT Imaging Facility and mouse genotyping platform (UMR9197 CNRS, NeuroPSI). We thank MIRCen’s and NeuroPSI’s animal facility and Chronobiotron (UAR3415) staff for mouse care. We thank Dr. Martine Cohen-Salmon and Dr. Katia Avila Gutierrez (CIRB, Collège de France) for their help with RNAscope experiments. We thank Nathalie Dechamps (Université Paris-Saclay, Inserm, CEA) for help with FACS experiments. We thank Dr. Jerrel Yakel (NIEHS, NIH, USA) and Dr. Uwe Maskos (Institut Pasteur) for respectively generating and providing α7nAChR floxed mice. We thank Dr. Alexis Faure and Dr. Julien Bouvier at NeuroPSI for providing ChAT-IRES-Cre and Ai9 mice, respectively.

This work was supported by the CNRS, the Université Paris-Saclay, a FRM “Retour en France” fellowship (ARF201909009244 to L.B.) and “Equipe” grant (EQU202303016285 to C.E. and L.B. and EQU202403018071 to Al.C.), a Fondation Vaincre Alzheimer pilot award (FR-20007p to L.B.), an ANR grants (ANR-24-CE16-0795-01 to L.B. and ANR-24-CE37-5566, ANR-23-CE37-0015-01 to Al.C.), a Centre National de la Recherche Scientifique and the Université de Strasbourg contract UPR3212 (to Al.C.), the NeuroStra Interdisciplinary Thematic Institute of the ITI 2021–2028 program (to Al.C.) and the ‘Fonds d’Investissement de l’INT jeunes chercheuses, jeunes chercheurs’ (FI_INT_JCJC_2021) (to R.B.). T.L and T.B received Ph.D. fellowships from Université Paris-Saclay (BIOSIGNE, 2021-2024) and Université Aix-Marseille (2021-2024).

## Author contributions

Designed the study: L.B., T.L., Performed surgeries: T.L., L.B., C.F., K-Y.W., M.G., C.E., An.C, Performed and analyzed behavioral experiments: T.L., L.B., C.F., M.G., S.B., Performed and analyzed fiber photometry experiments: C.F., Performed histological and biochemical experiments: T.L., L.B., C.F., C.J., S.B., Performed microscopy and analysis of histological data: T.L., L.B., C.F., Performed and analyzed electrophysiological experiments: T.B., R.B., K-Y.W., V.G., Training and lab management: M-A.CS., Wrote first version of the article: L.B., Reviewed and edited article: All co-authors, Acquired funding: L.B., C.E, Al.C., R.B. All authors approved the manuscript.

---

**Extended data Fig.1:**
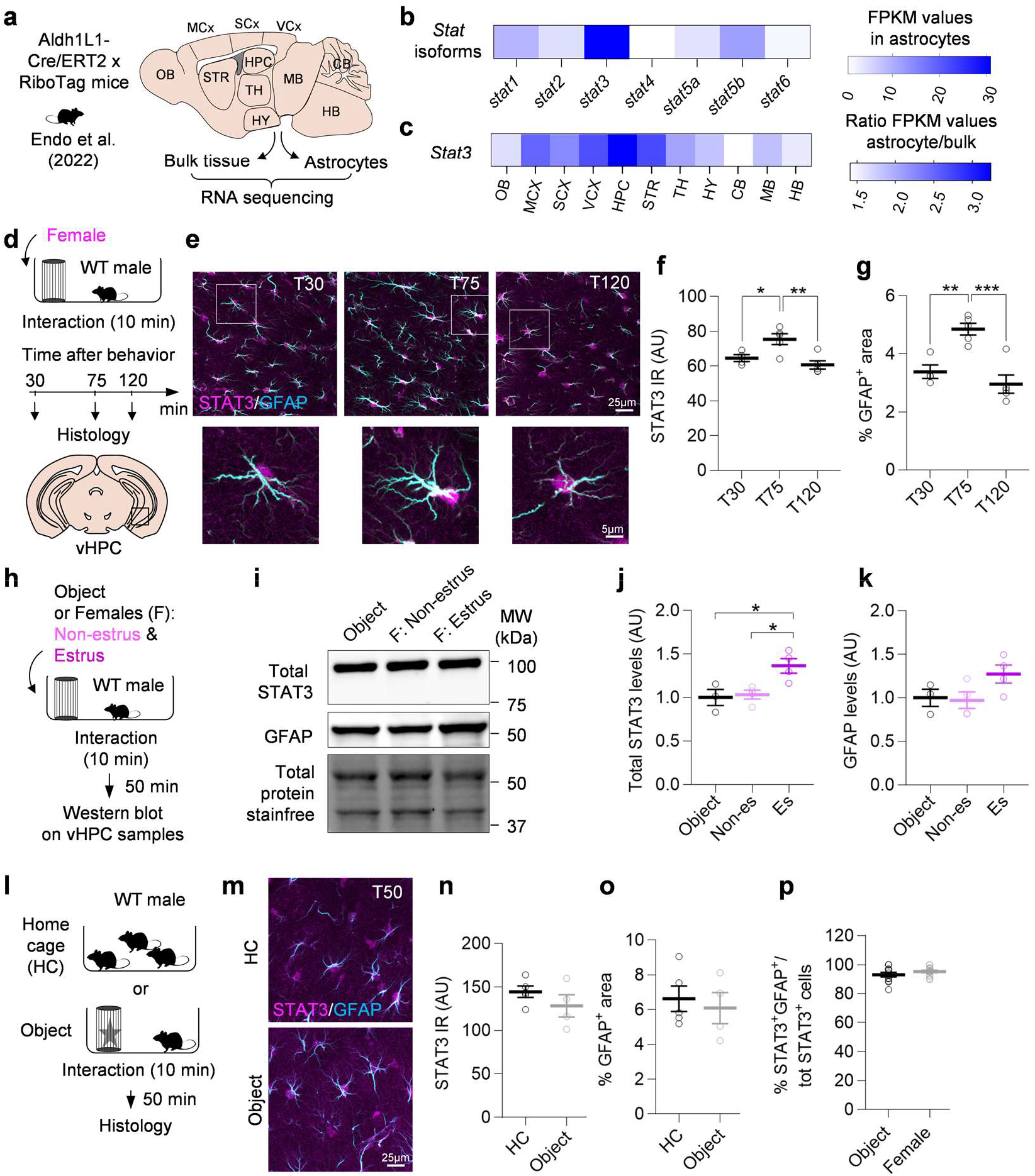
STAT3 is transiently and specifically induced in vHPC astrocytes of male mice after exposure to a female. **a,** Experimental design used by Endo et al. 2022. *Aldh1l1-cre^ERT2^* mice were bred with RiboTag mice to induce the expression of *Rpl22-HA* in astrocytes upon tamoxifen injection allowing the immunoprecipitation of astrocyte-specific ribosome-bound mRNAs from 11 brain regions. RNA was either extracted and sequenced from bulk tissue lysates or from the immunoprecipitation samples containing astrocyte-enriched RNA ^29^. **b,** FPKM values of *Stat* isoforms. **c,** Ratio of *Stat3* FPKM values in astrocytes versus bulk tissue in 11 brain regions. Abbreviations: OB: olfactory bulb, MCX: motor cortex, SCX: Somatosensory cortex, VCX: visual cortex, HPC: hippocampus, STR: striatum, TH: thalamus, HY: hypothalamus, CB: cerebellum, MB: midbrain, HB: hindbrain. **d,** WT male mice were exposed to an adult female for 10 min and euthanized 30 min, 75 min or 120 min later. **e**, Confocal images of vHPC sections from male mice at different time points after exposure to a female, co-stained for STAT3 (magenta) and GFAP (cyan). **f, g,** Quantification of nucleo-somatic STAT3 immunoreactivity (IR) (**f**) and the % GFAP+ image area (**g**) at different time points (n=4-5 mice/group). **h**, Adult WT male mice were let to interact either with an object (Obj) or non-estrus (Non-es) or estrus (Es) females. After 50 min, vHPC samples of male mice were collected and processed for western blotting. **i**, Representative western blotting images of STAT3, GFAP and total proteins detected on StainFree membrane. **j, k**, Quantification of STAT3 (**j**) and GFAP (**k**) protein levels normalized by the total protein amount (n=3-4 mice/group). **l**, WT male mice were either let to interact with an object for 10 min or stayed in their home cage. After 50 min, their brains were processed for histology. **m,** Confocal images of vHPC sections co-stained for STAT3 (magenta) and GFAP (cyan), from mice left in their home cage or after a 10-min interaction with an object. **n, o**, Quantification of STAT3 IR (**n**) and % GFAP+ area (**o**) in home cage and object conditions (n=4-5 mice/group). **p**, Quantification of the proportion of STAT3+ cells also expressing GFAP, after interaction with an object and a female (n=12-15 mice/group). **f, g, j, k**: One-way ANOVA and Tukey’s multiple comparison tests; **n**, **o**, **p**: Two-tailed unpaired t-test. *, p<0.05, **, p<0.01, ***, p<0.001. Values were plotted as mean ±SEM.

**Extended data Fig.2:**
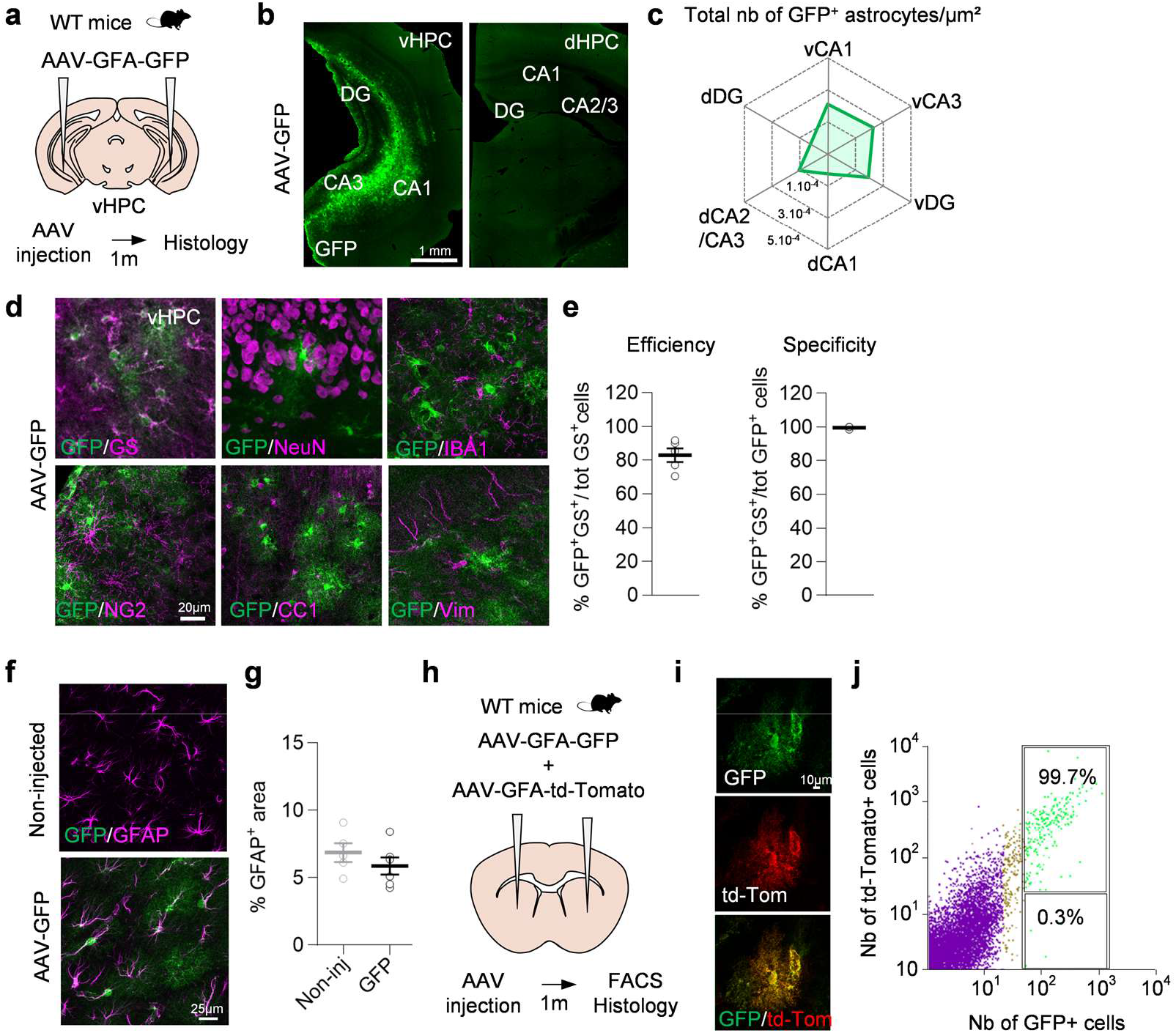
Characterization of astrocyte-targeted AAV diffusion and tropism in the mouse vHPC. **a**, Adult WT mice received bilateral injections in the vHPC of astrocyte-targeted AAV encoding GFP. **b,** Low magnification images of GFP immunostaining on vHPC (left) and dHPC (right) brain sections from AAV-GFP-injected mice. **c,** Spider plot representing the number of GFP+ astrocytes per µm² in different HPC subregions (n=15 mice). **d,** Confocal images of brain sections from AAV-GFP injected mice co-stained with cell type-specific markers (magenta) for astrocytes (GS), neurons (NeuN), microglia (IBA1), oligodendrocyte progenitor cells (NG2), mature oligodendrocytes (CC1) and radial glia-like cells (Vim). **e**, Quantification the transduction efficiency and specificity. **f**, Confocal images of the vHPC of non-injected and AAV-GFP-injected mice co-stained for GFP and GFAP. **g**, Quantification of the % GFAP+ image area (n=5-6 mice/group). **h**, Adult WT mice received intracerebral injections of a mix of astrocyte-targeted AAV encoding GFP and td-Tomato. Brains were processed for histology or the injected area was dissected out for FACS analysis. **i**, Confocal image of a co-transduced cell expressing both GFP and td-Tomato. **j**, Percentage of cells expressing GFP, td-Tomato or both was evaluated based on their fluorescence by FACS (n=1 sample pooled from 4 mice). **g**: Two-tailed unpaired t-test. Values were plotted as mean ±SEM.

**Extended data Fig.3:**
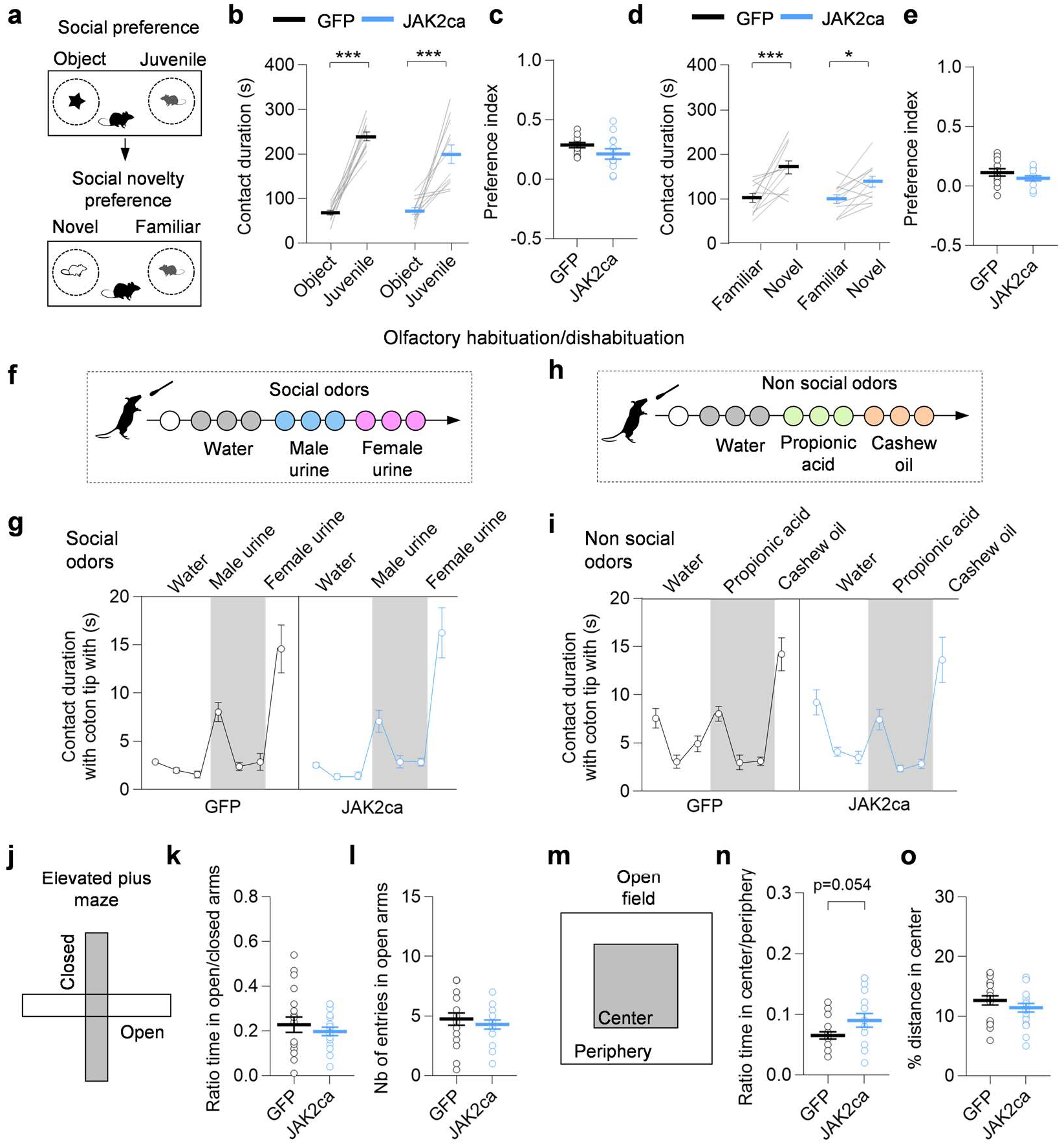
Experimental activation of astrocyte JAK2-STAT3 signaling in the vHPC selectively impacts male behavior towards females. **a,** WT male mice injected in the vHPC with astrocyte-targeted AAV encoding J*ak2ca + Gfp* or *Gfp* were tested to assess their sociability and social novelty preference towards juveniles. **b, c**, Contact duration (**b**) and preference index (**c**) for the social preference phase (n=12 mice/group). **d, e**, Contact duration (**d**) and preference index (**e**) for the social novelty preference phase (n=12 mice/group). **f-i**, Mice were tested at the olfactory habituation/dishabituation test and the contact duration with social (**f, g**) and non-social (**h, i**) odors was determined (n=16-19 mice/group). **j**, AAV-JAK2ca and GFP-injected males were tested at the elevated plus maze to evaluate anxiety-like behaviors. **k, l,** Ratio of time spent in the open versus the closed arms (**k**) and numbers of entries in open arms (**l**) at the elevated plus maze between GFP and JAK2ca mice (n=18-19 mice/group). **m**, AAV-JAK2ca and GFP-injected males were tested at the open field to evaluate anxiety-like behavior. **n, o**, Ratio of time spent in the center versus periphery (**n**) and % of distance spent in the center (**o**) are compared between groups (n=18-20 mice/group). **b**, **d, g, i**: Two-way repeated measure ANOVA and Sidak’s multiple comparison tests; **c**, **e**, **k, l**, **n**, **o**: Two-tailed unpaired t-test. *, p<0.05; ***, p<0.001. Values were plotted as mean ±SEM.

**Extended data Fig.4:**
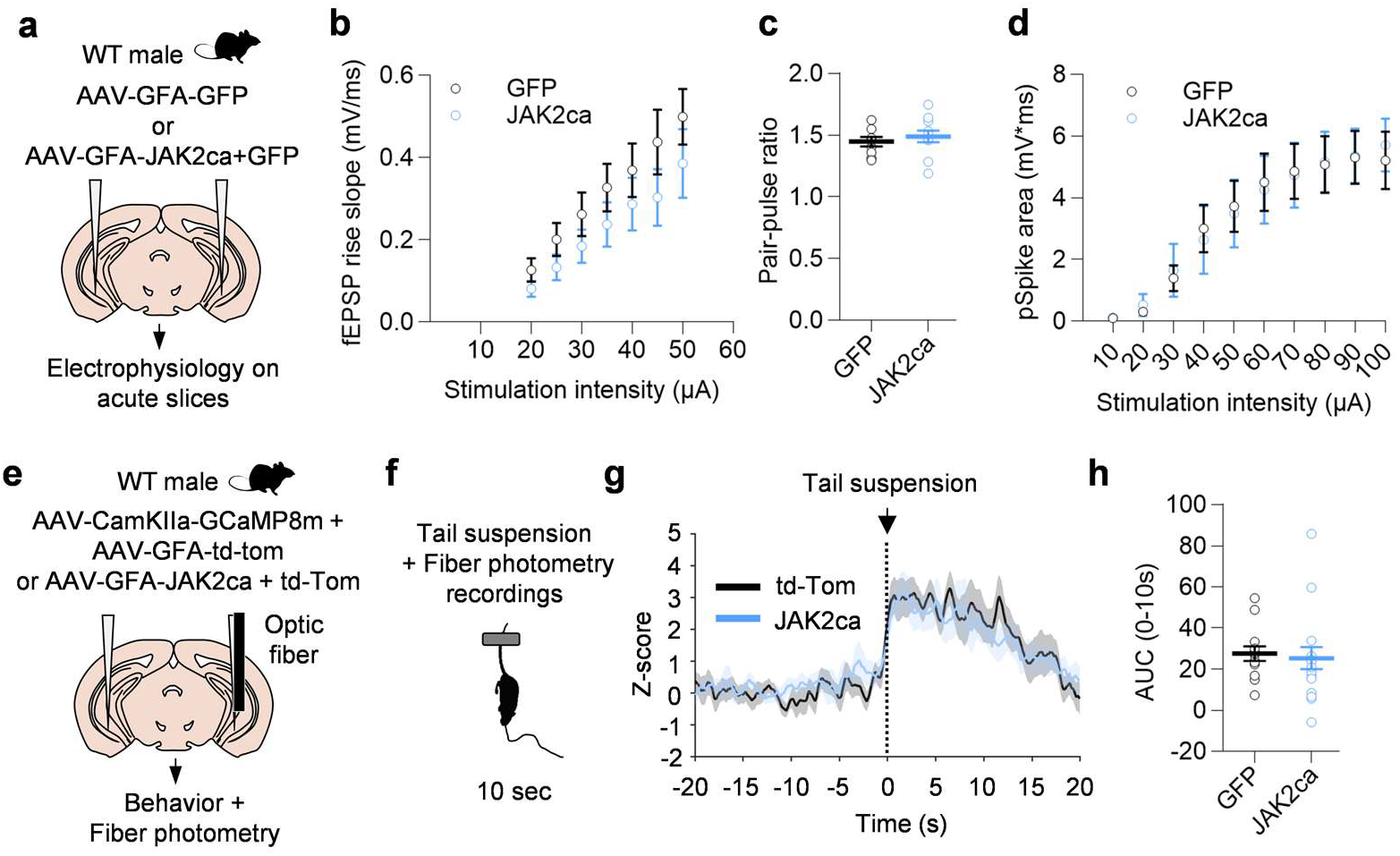
Experimental activation of astrocyte JAK2-STAT3 signaling in the vHPC does not impact pyramidal neuron response to tail suspension and AMPAR activity at the Schaffer’s collateral-vCA1 synapse. **a,** Field excitatory post-synaptic potentials (fEPSPs) recordings were performed on acute vHPC slices of AAV-JAK2ca and -GFP-injected mice. **b**-**d**, fEPSP rise slope (**b**), paired-pulse ratio (**c**) and spike population (**d**) plots are represented (n= 3-7 slices from 4 mice/group). **e, f**, Adult WT males received bilateral intracerebral injections of a mix of AAVs targeting neurons and encoding the genetic Ca2+ indicator (GCaMP8m) and of AAVs targeting astrocytes enabling the expression of JAK2ca and/or td-Tomato. Mice were implanted in the vHPC with an optic fiber for subsequent fiber photometry recordings during tail suspension. **g, h**, Z-score (**g**) and AUC (**h**) plots of vCA1 neurons Ca2+ activity for the tail suspension (n= 13-18 mice/group). **b**, **d**: Two-way repeated measure ANOVA and Sidak multiple comparison test; **c**: Two-tailed unpaired t-test; **g**: Two-sided permutation test; **h**: Mann Whitney test. Values were plotted as mean ±SEM.

**Extended data Fig.5:**
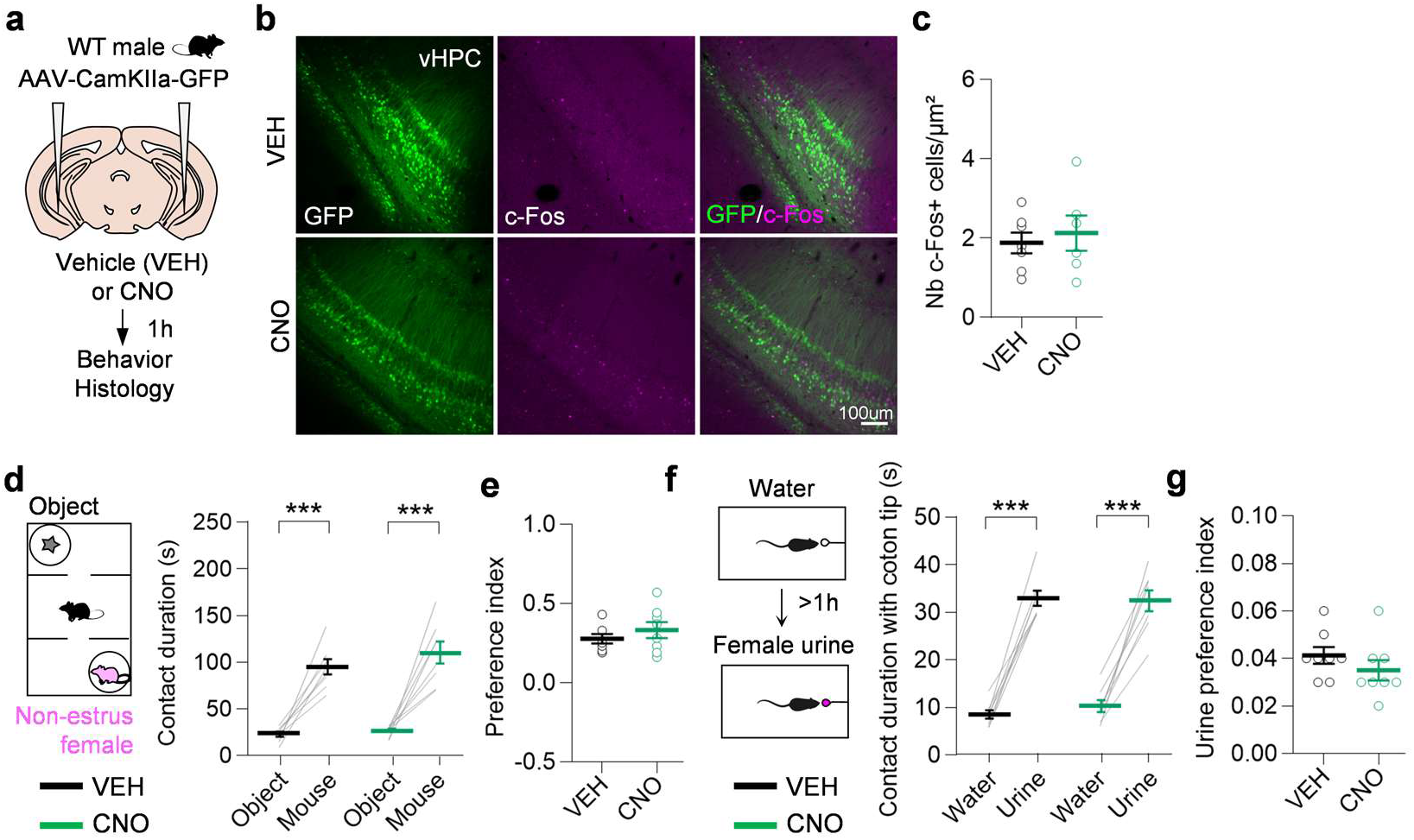
Systemic CNO administration in non-hM3Dq-expressing mice does not lead to prominent behavioral changes. **a,** WT male mice received intracerebral injections in the vHPC of AAVs targeting pyramidal neurons (CaMKIIa promoter) and encoding a control transgene, GFP. One-month post-injection, mice were administered with CNO (0.1 mg/kg) or vehicle (VEH) via micropipette-assisted drug administration one hour prior to histological analysis or behavioral testing. **b**, Confocal images of vHPC brain sections from AAV-CamKIIa-GFP-injected mice after administration of VEH or CNO stained for GFP (green) and c-Fos (magenta). **c**, Quantification of number of c-Fos+ cells in the transduced area (n=6-7 mice/group). **d**, **e**, Contact duration with object and non-estrus females (**d**) and preference index (**e**) at the three-chamber test (n=8 mice/group). **f, g**, Contact duration with water and female urine (**f**) and preference index (**g**) at the female urine sniffing test (n=8 mice/group). **c**, **e**, **g**: Two-tailed unpaired t-test; **d**, **f**: Two-way repeated measure ANOVA and Sidak’s multiple comparison tests. ***p < 0.001. Values were plotted as mean ±SEM.

**Extended data Fig.6:**
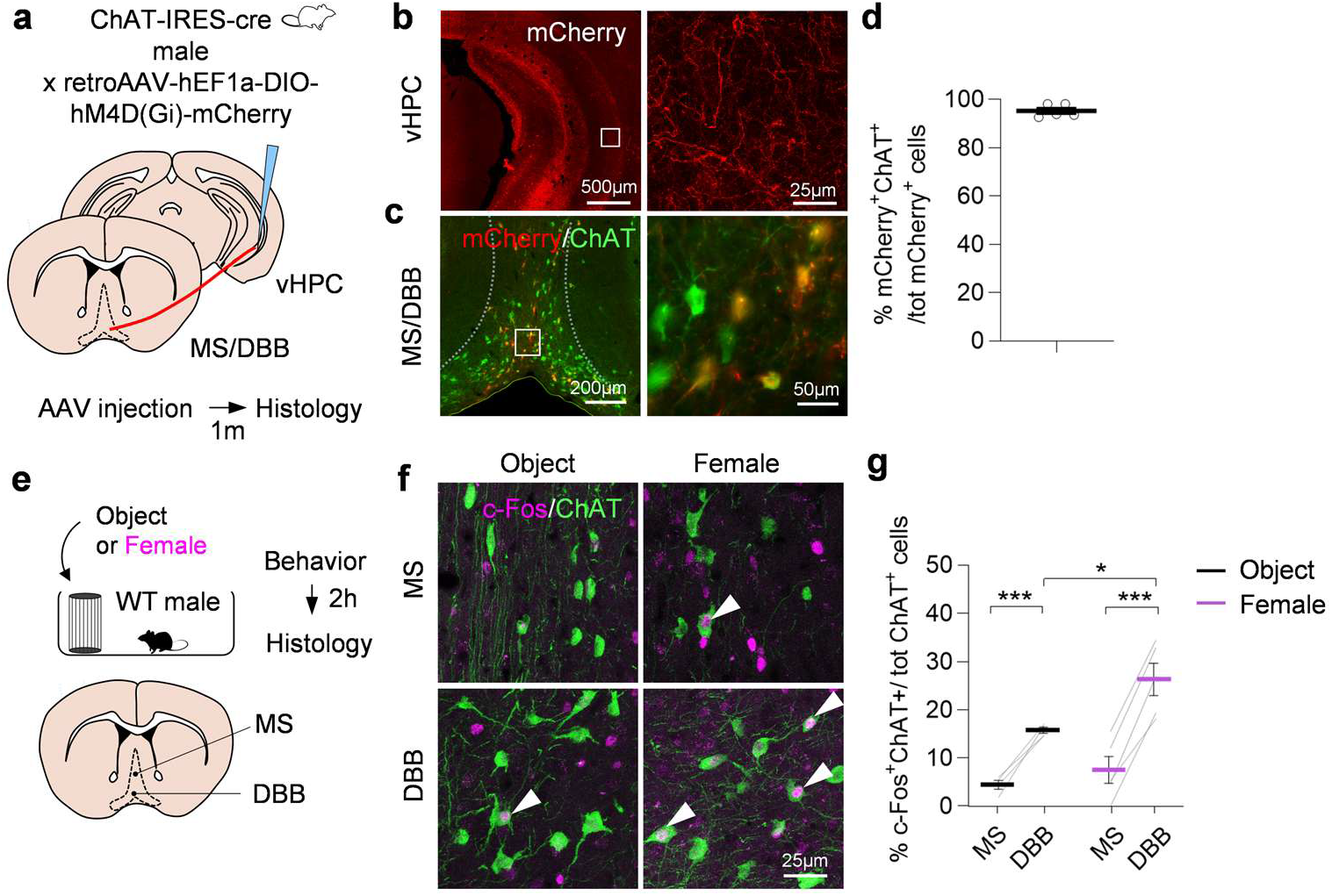
The mouse vHPC receives cholinergic inputs from the MS/DBB, which are differentially activated following female exposure. **a,** ChAT-IRES-cre male mice received intracerebral injections of a Cre-dependent retrograde AAV in the vHPC encoding the inhibitory DREADD, hM4Di-mCherry. One month later, their brains were analyzed by histology. **b**, Low (left) and high (right) magnification images showing the viral-mediated mCherry expression in vHPC-projecting axons (red). **c,** Retrogradely-labeled mCherry+ vHPC-projecting neurons (red) in the MS/DBB express the cholinergic neuron marker ChAT (green). **d**, Quantification of the % of mCherry+ChAT+ neurons over the total number of ChAT+ neurons in the MS/DBB (n=4-5 mice/group). **e**, WT male mice were let to interact with an object or an estrus female for 10 min. Two hours later, mice were perfused and their brains analyzed by histology. **f**, Confocal images of MS and DBB brain sections of male mice after interaction with an object or a female and co-stained for ChAT (green) and c-Fos (magenta). **g**, Quantification of the % of c-Fos+ChAT+/total number of ChAT+ neurons in the MS and the DBB (n=4-5 mice/group). **g**: Two-way repeated measure ANOVA and Sidak’s multiple comparison tests. *, p<0.05, ***, p<0.001. Values were plotted as mean ±SEM.

**Extended data Fig.7:**
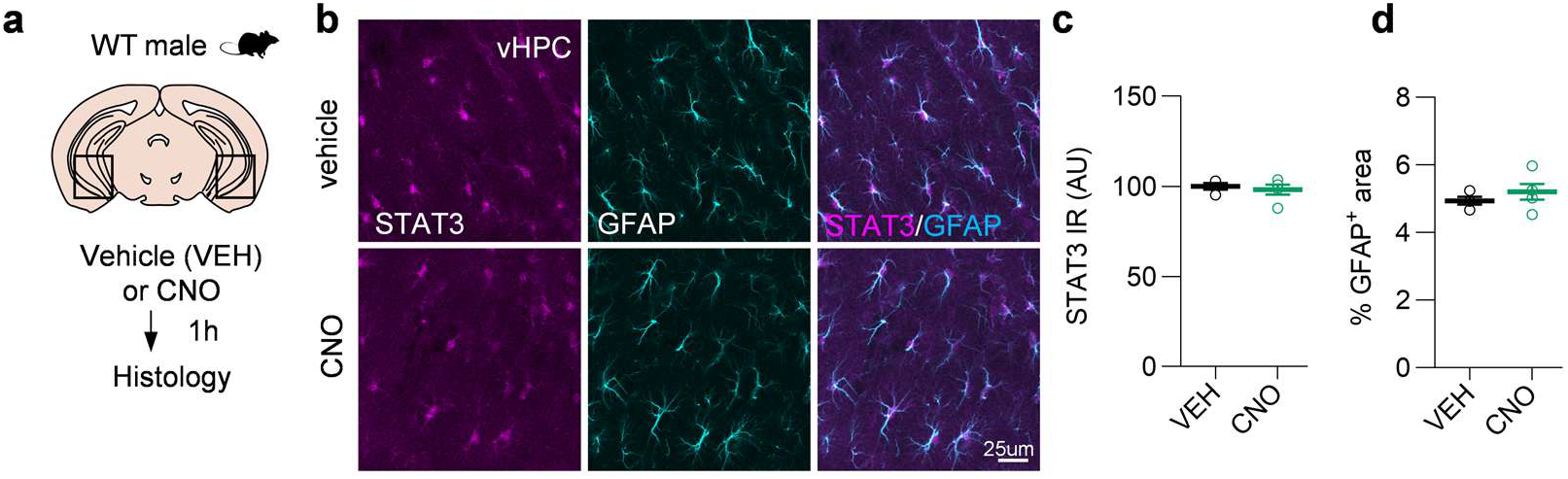
Systemic administration of CNO does not activate the JAK-STAT3 signaling in vHPC astrocytes. **a,** WT male mice were administered with vehicle (VEH) or CNO (2 mg/kg) 1h prior to perfusion for brain analysis by histology. **b,** Confocal images of vHPC brain sections mice administered with VEH and CNO, co-stained for STAT3 (magenta) and GFAP (cyan). **c, d**, Quantification of nucleo-somatic STAT3 immunoreactivity (**c**) and the % GFAP+ image area (**d**) in VEH and CNO mice (n=4-5 mice/group). **c, d**: Two-tailed unpaired t-test. Values were plotted as mean ±SEM.

**Extended data Fig.8:**
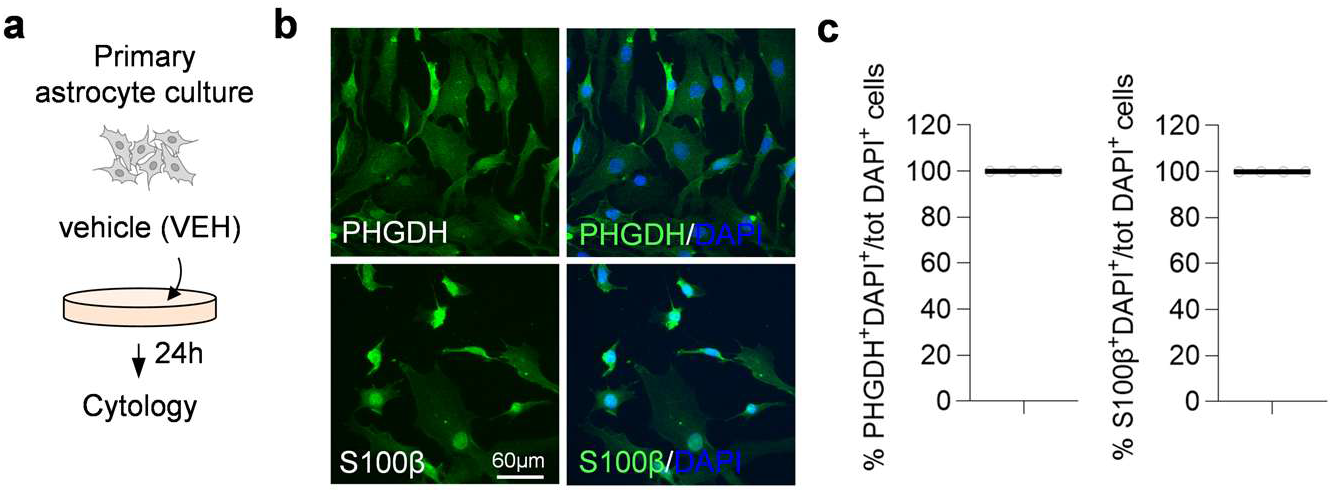
Validation of primary murine astrocyte culture purity. **a,** Control primary astrocyte cultures were treated with DMSO (VEH) and processed for immunocytology after 24h. **b**, Epifluorescence images of primary astrocyte cultures co-stained with astrocyte-specific markers PHGDH and S100b (green) and DAPI (blue). **c**, Quantification of the proportion of PHGDH+ and S100b+ cells over the total number DAPI+ of cells (n=4 wells from 2 cultures). Values were plotted as mean ±SEM.

**Extended data Fig.9:**
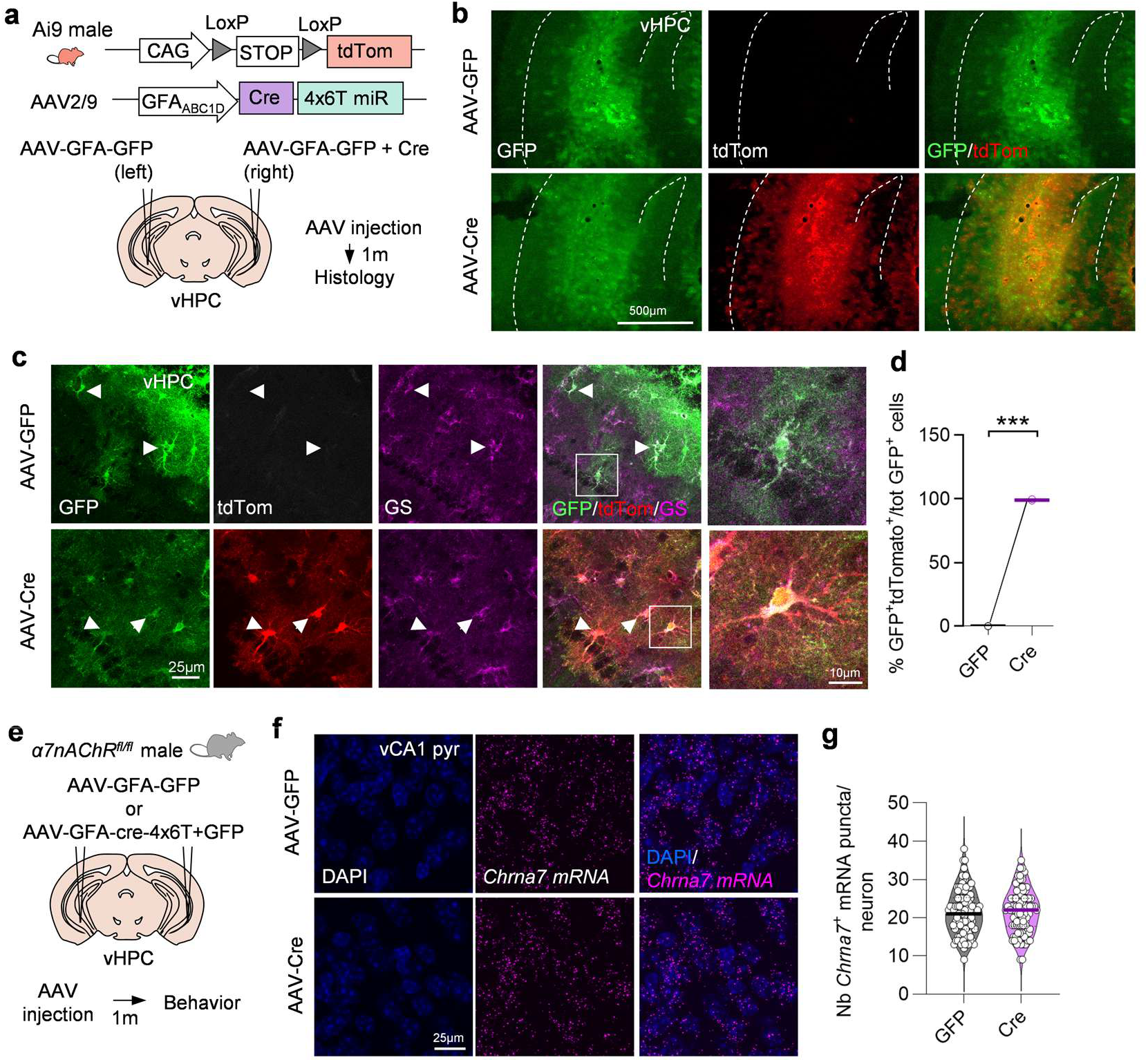
Validation of astrocyte-specific α7nAChR cKO strategy in in the mouse vHPC. **a,** Ai9 td-Tomato Cre reporter mice were injected in the vHPC with an AAV targeting astrocytes and encoding *Cre* mixed with AAV-GFP (right hemisphere) and AAV-GFP control (left hemisphere). Mouse brains were analyzed by histology 1 month after viral injections. **b,** Epifluorescence images of vHPC brain sections injected with AAV-GFP (top) and AAV-cre (bottom) following immunofluorescent amplification of virally-encoded GFP and td-Tomato. **c**, Confocal images of vHPC brain sections of AAV-Cre and -GFP-injected mice co-stained for GFP (green), td-Tomato (red) and the astrocyte marker GS (magenta). **d**, Quantification of the % of GFP+td-Tomato+/total GFP+ astrocytes in the GFP and Cre groups (n=4 mice/group). **e,** Adult α7nAChR^fl/fl^ mice received bilateral injections in the vHPC of AAV targeting astrocytes and encoding either a control *Gfp* or *Cre + Gfp*. One month after injection, mice underwent behavioral analysis. **f,** Confocal images of vHPC CA1 pyramidal layer identified by DAPI and co-labelled by fluorescent *in situ* hybridization for *Chrna7* mRNA. **g**, Quantification of the number of *Chrna7+* puncta per DAPI+ pyramidal neuron in the AAV-GFP and -Cre groups (n=87-106-cells from 3 mice). **d**: Two-tailed paired t-test; **g**: Linear mixed model (fixed effect: group; random effect: mouse). ***, p<0.001. Values were plotted as mean ±SEM.

## Materials and Methods

### Mice

Adult male C57BL6/JRj (Janvier labs), Aldh1L1-GFP (GENSAT), homozygous ChAT-IRES-Cre (ChAT-IRES-Cre::SV40pA::frt-neo-frt, JAX #006410), homozygous α7nAChRfl/fl (provided by Jerrel Yakel, NIH and Uwe Maskos, Institut Pasteur) and homozygous Ai9 mice (B6. Cg-Gt (ROSA)26Sortm9(CAG-tdTomato) Hze/J, JAX #007909) male mice of 3-6 months were used as experimental subjects. For genetic labeling of cholinergic neurons, ChAT-IRES-Cre male mice were crossed with Ai9 females. Juvenile males or females (1.5-2 months) and adult females (3-6 months) C57BL6/J were used as stimuli for social interaction assays and born either in our animal facility or purchased (Janvier labs). Animals were housed under standard conditions (12h light-dark cycle, temperature: 22 ± 1 °C and humidity: 50%) with *ad libitum* access to food and water. All experimental protocols were reviewed and approved by the local ethics committee (CEEA N°44, 59, 35 and 71) and submitted to the French Ministry of Education and Research (#24301-202002241344372 v1, #34228-2021111710469619 v12, #17485-2018110819197361 and #44572-2023072115506121). They were performed in authorized facilities (#B92-032-02 and #E91-272-108), in strict accordance with recommendations of the European Union (2010–63/EEC). All behavioral experiments were performed during the light phase, between 8 am and 2 pm under dim light (∼30-50 lux) unless stated otherwise. Experimental subjects were group housed by up to 5 and sexually naïve prior to behavioral testing. To avoid any potential cage effect, mice injected with control and test AAVs were always co-housed in varying proportions. Experimenters were blind to group when performing behavioral testing and video annotation. When mice were euthanized shortly after behavioral testing (50 min-2h), conditions were randomized to avoid time-of-the-day effects. When possible, mice from the same cage were exposed to the same type of stimulus (i.e. object or female) and put back together in their home cage to avoid cross-contamination of social signals between individuals.

### Viral vectors

To target astrocytes, mice were injected with AAVs of serotype 2/9 and 2/5 carrying a synthetic promoter derived from the GFAP promoter (gfaABC1D) ^45^ and detargeting sequences (miR) to prevent potential ectopic expression in neurons ^35,46^. We used viral vectors encoding either murine *Jak2^T875N^*(a constitutively active form of JAK2 (JAK2ca)) and *Socs3* (the endogenous pathway’s inhibitor) to respectively activate and inhibit the JAK2-STAT3 signaling ^16^, a *Cre* recombinase (Addgene, #196410) or fluorescent proteins, *Gfp* and *td-tomato*. To target vHPC pyramidal neurons, we used AAVs encoding *hM3Dq-mCherry* (Addgene #50476), *Gcamp8m* (Addgene, #176751) and *Gfp*, under the control of a *CamkIIa* promoter. For the chemogenetic inhibition of cholinergic inputs onto the vHPC, we used a Cre-dependent (double floxed inverted open reading frame, DIO) retrograde AAV encoding *hM4Di-mCherry* under the ubiquitous *eIF1a* promoter (Brain VTA, #PTA-0043). AAVs were either obtained from the Molecular Imaging Research Center (MIRCen) viral core, Addgene or Brain VTA. Stock AAV solutions were diluted in 0.1 M PBS/0.001% pluronic acid unless specified otherwise. Retro-AAV-DIO-eIF1a-hM4Di-mCherry were injected undiluted at 5.6.10⁹ viral genome (VG)/site. AAV2/9-CaMKIIa-hM3Dq-mCherry and -GFP were injected at 5-6.10⁹ VG/site. Mice injected with astrocyte-targeted AAVs received a total of 5.10⁹ VG/site. To visualize transduced cells in groups where the transgenes cannot be detected by histology due to a lack of efficient antibodies (i.e SOCS3 and JAK2), we used a mix of AAVs targeting astrocytes at a 1:4 ratio with a reporter AAV encoding a fluorescent protein (GFP or td-Tomato). For the same reason, when mixing astrocyte- and neuron-targeted AAVs, mice received a total load of 8.10⁹ VG/site, containing 3.10⁹ VG of neuronal and 5.10⁹ VG of astrocyte AAV.

### Chemogenetic manipulations

Clozapine-N-Oxide (CNO, Enzo, #BML-NS105) was dissolved in water at 6 mg/mL, aliquoted and stored at −20°C. On the day of the experiment, mice received CNO at 0.1 mg/kg (for activation) and 2 mg/kg (for inhibition) or vehicle (water) in 40% sweetened condensed milk solution through non-invasive micropipette-guided drug administration (MDA) ^47^1h before behavioral testing or before euthanasia for subsequent histological analysis. Briefly, the week before MDA habituation, diluted condensed milk was first pipetted onto the house cage grid of mice to limit neophobia. Then, mice were successively habituated to drink the condensed milk without CNO from a P200 pipette tip while on the house cage’s grid first using contention (day 1), then only a slight tail grab (days 2 and 3) and finally voluntarily (day 4). A small proportion of mice that were still refractory to micropipette administration were grabbed by the tail on the test of the experiment.

### Stereotaxic surgeries

Mice were anesthetized either using a subcutaneous injection of a mix of ketamine (100 mg/kg) and medetomidine (0.25 mg/kg) or with 4% isoflurane in an induction chamber and maintained with 1.5-2% isoflurane during the surgery. Analgesia was achieved by a subcutaneous injection of xylocaine (5 mg/kg) at the incision site and of buprenorphine (0.075 mg/kg) or 5% lidocaine cream was applied on the scalp and local analgesics (Bupivacaine, 2 mg/kg, and Lidocaine, 2 mg/kg) were injected locally at the incision site 30 min before the end of the surgery. Metacam (Meloxicam, 10 mg/kg) was injected subcutaneously (s.c.) to limit inflammation before surgery) for 2 days after surgery. Corneal protection gel (Ocrygel) was applied to both eyes before placing mice onto the stereotaxic frame. Mouse body temperature was monitored and maintained starting from the beginning of anesthesia to the recovery using temperature-controlled heating pads and recovery chamber. Viral vectors were injected bilaterally in the vHPC using the following coordinates based on the Allen Brain Atlas relative to Bregma (antero-posterior (AP): −3.0 mm, medio-lateral (ML): ±3.5 mm, dorso-ventral (DV): −4.5 mm) in a total volume of 1-2 µl, at a rate of 0.25 µl/min. Once the injection completed, injection cannulas were left in place for 5 min to facilitate AAV diffusion before removal. For fiber photometry, following viral injections during the same surgery, optic fiber (400 µm diameter, 0.39 NA, RWD, R-FOC-BL400C-39NA) were implanted on the right side, 100 µm above the DV coordinate used for viral injection. A screw was superficially inserted onto the skull and secured with the cannula using adhesive dental cement (Superbond (Phymep, #K058E) and Paladur (Kulzer, #64707965). At the end of surgery, wounds clips or sutures were used to close the incision, pre-warmed saline (10 ml/kg) was injected subcutaneously in the flanks for rehydration. Anesthesia recovery time was reduced by a subcutaneous injection of the reversal agent Atipamezole (0.25 mg/kg). Mice were let to recover in a heated chamber and returned to their home cage once they resumed walking. Wounds clips or sutures were removed 10 days after surgery under isoflurane anesthesia (4% induction in medical air) while mouse habituation prior to behavioral testing was performed at least 1-month post-surgery to allow mice full recovery and transgene expression.

### Behavioral analysis

#### Habituation and data acquisition

Prior to behavioral testing, mice were handled by the experimenter (2 min/mouse for 3-5 consecutive days). Stimuli mice were habituated to the barred cage (5-10 min/mouse for 3 days). One stimulus mouse was exposed to maximum 4 test mice, non-consecutively. The mouse geometric center was detected and video-tracked using EthoVision XT (v15-17). Nose or paw contacts through the barred cage were either live- or *a posteriori* annotated on videos. All behavioral testing apparatuses were thoroughly cleaned with 10% ethanol between each mouse to avoid inter-individual olfactory contamination. Expression of viral-mediated transgene expression was confirmed by histology when possible and mice with no fluorescent cells or incorrect targeting were removed from all analyses.

#### Estrus cycle evaluation

Vaginal cytology was performed as described in Mc Lean et al.^48^. Soiled litter from a cage of male mice was systematically added to the female cages at least one week prior to behavioral testing to increase cycle synchronicity. Female mice were restrained and ∼50 µl of saline was gently injected into the vaginal cavity to collect cells, which were then spread onto glass slides, air-dried and stained with Crystal violet. Cycle phase was determined following microscopic evaluation of cellular morphology. To avoid interference of restrain-induced stress in females with behavioral testing and disruption of the vaginal advertising molecules, the estrus cycle evaluation was performed 1-4 days prior to testing and an estimation was calculated ^49^. When possible, cycle was confirmed with post-testing swab. Only females with the intended estrus cycle phase were subsequently used for behavioral assays.

#### Social and non-social stimulus exposure

Test male mice were put in an empty house cage with the barred cage containing either an object (15 ml Falcon tube filled with colorful content) or an adult female. Test mice were let to freely explore for 10 min, then returned to their home cage and euthanized 60 min later.

### Social preference test

Mice were placed in an empty three-chamber apparatus (60 x 40 x 22 cm, Ugo Basile) and allowed to freely explore for 10 min. They were then gently returned to the central chamber with closed doors while an object and a female were placed in the barred cages in opposite chambers of the apparatus. After the doors were reopened, mice were allowed to interact with the stimuli for 5 min. The time spent in contact (sniffing or grabbing with forepaws) with the barred cages were quantified. A preference index was calculated as (time in contact with female – time in contact with object)/total duration. Position of the object and social stimuli was alternated to avoid place preference. Z-score were calculated on contact duration with females for each animal as (contact duration in test - mean (contact duration in controls)/standard deviation (contact duration in controls)).

#### Social novelty preference test

Behavioral testing was conducted in a single arena (20 x 40 x 22 cm), with two barred cages placed at the extremities. After a habituation phase of 10 min with empty containers, mice were let to explore the two barred cages each containing an object or a juvenile (familiar) male for 10 min. Next, the object was replaced by another juvenile (novel) male and the test mice let to explore for another 10 min. The contact (sniffing or grabbing with forepaws) with the barred cages were quantified. A preference index was calculated as (time in contact with novel mouse – time in contact with familiar mouse)/total duration.

#### Female Urine Sniffing Test

Test male mice were placed in a house cage with clean litter and a cotton swab secured onto one wall, 8 cm above the litter so that mice can reach it by rearing. During the habituation phase, water (18 µl) was pipetted onto the cotton swab while urine was applied during the test phase. For each phase, mice were let to freely explore the cage for 5 min with an inter-trial interval of at least 45 min. The cumulative sniffing duration was manually scored using EthoVision XT. Urine was collected in adult C57BL6/J males and females during 5 days the week preceding the test and instantly frozen on dry ice. A master mix containing the same volume of urine from each animal was prepared, aliquoted and stored at −20°C until use so that each test mouse was exposed to the exact same urine sample. A preference index for the female urine was calculated as (contact duration with urine – contact duration with water)/ total duration.

#### Olfactory habituation/dishabituation test

Mice with normal olfaction display a decrease in contact duration when consecutively exposed to the same odor (olfactory habituation) and an increase in sniffing when exposed to a novel odor (dishabituation). We performed this test first with non-social odors (water (3 times), diluted propionic acid (1% (v/v), 3 times) and cashew oil (10% (w/v), 1 time)), then one week later with social odors (water (3 times), female urine (3 times) and male urine (1 time)). After 15 min habituation in a housing cage with clean litter, mice were allowed to contact a cotton swab, secured on one cage wall for 3 min with an inter-trial interval of 2 min. The cumulative sniffing duration was manually scored.

#### Elevated Plus Maze test

Test mice were placed in an elevated plus-shaped maze (arm length: 35 cm, arm width: 5 cm, height: 61 cm) with two open arms (150 lux) perpendicular to two closed arms (30 lux) for 6 minutes. Time spent in open and closed arms and number of entries in open arms were automatically calculated using EthovisionXT.

#### Open field test

Mice were placed in an open field arena (60 x 60 x 40 cm) and let to freely explore for 10 minutes. Total distance moved and cumulative time spent in the center and in the periphery of the arena were automatically calculated using EthovisionXT.

### Fiber photometry recordings

Prior to any behavioral testing, mice were handled by the experimenter for 2 days and then habituated to the weight and feel of the optic fiber cable while placed in a house cage for 5 min for 2 consecutive days. On test day, mice were connected to the patchcord and then placed in a house cage with an empty barred cage for at least 1min before recording started. Signals were acquired using a commercial multichannel fiber photometry system and software with two light-emitting LEDs: 410 nm (isobestic signal) and 470 nm (GCaMP signal) (RWD life science). The signal was acquired during 5 min without any stimulus in the cage, then an object was placed in the barred cage and left in place for 2 min, before being removed for 1 min and being presented again twice. After the object presentation, a non-estrus female was placed in the barred cage and left in place for 2 min. Behavioral video files and fluorescence data were acquired at 60 Hz. Plots were centered to the first contact with the barred cage. Fiber photometry data were analyzed using custom scripts written in MATLAB (MathWorks, Natick, MA, USA) and available in the following GitHub repository: https://github.com/LucileBH/Lakomy_et_al._2026. Fluorescence signals acquired at 410 nm and 470 nm were imported into MATLAB for preprocessing and analysis. The first minute of recording was excluded from all analyses to avoid potential instability associated with the initial recording period. <u>Signal preprocessing:</u> Both fluorescence channels were first denoised using a median filter and were then low-pass filtered (fourth-order Butterworth filter, cutoff: 10 Hz). <u>Photobleaching correction:</u> Photobleaching was corrected independently for the 410 nm and 470 nm channels by fitting each trace with a double-exponential decay model. The fitted photobleaching curve was then subtracted from the filtered fluorescence trace to obtain photobleaching-corrected signals. <u>Motion artifact correction:</u> To remove movement-related artifacts, a linear regression was performed between the corrected isosbestic (410 nm) and Ca2+-dependent (470 nm) signals. The predicted motion artifact was calculated from the fitted regression and subtracted from the corrected 470 nm signal. <u>Peri-event analysis, z-score normalization and AUC:</u> Peri-event fluorescence responses were normalized to the median fluorescence during the interval from −15 s to −5 s preceding the contact with the barred cage (baseline median fluorescence, F0). Event-related fluorescence traces were normalized using z-score transformation: z-score = (F(t)-F0)/ σF0. The AUC was calculated using the trapezoidal numerical integration method (trapz function in MATLAB), providing an estimate of the integrated signal over time. Three animals were excluded based on the correlation between the 410 nm isosbestic and 470 nm Ca2+-dependent signals. Two animals exhibited a negative correlation coefficient (410/470 correlation < 0), indicative of abnormal signal dynamics that were accompanied by aberrant behavior consistent with seizure-like activity. One additional animal displayed a high correlation coefficient (r = 0.90), suggesting that the recorded fluorescence was predominantly driven by background fluctuations rather than Ca2+-dependent activity.

### Electrophysiological recordings

#### Slice preparation

Animals underwent cervical dislocation and decapitation. For patch-clamp recordings, mice were deeply anesthetized with ketamine/xylazine (100/10 mg/kg, i.p.) and transcardially perfused with an ice-cold N-methyl D-glucamine (NMDG)-based solution containing (in mM): 93 NMDG, 2.5 KCl, 1.2 NaH2PO4, 30 NaHCO3, 20 HEPES, 20 glucose, 10 MgCl2, 93 HCl, 2 Thiourea, 3 sodium pyruvate, 12 N-acetyl cysteine, and 0.5 CaCl2 (equilibrated with 95% O2 and 5% CO2, pH 7.2– 7.4). The brain was then removed from the skull and both hippocampi were dissected, aligned longitudinally and introduced into a 1% agar solution, quickly cooled, fixed in an agar block (2.25% weight/volume in aCSF). The agar block containing both hippocampi is then mounted in a vibrating microtome (Leica, VT1000S) and sliced (350 μm) in ice-cold oxygenated NMDG-based solution. Slices were immediately transferred to recover in NMDG-based solution at 35°C for 5 min and then stored for at least 1 h at room temperature in normal artificial cerebrospinal fluid (ACSF) containing (in mM): 126 NaCl, 2.5 KCl, 1.2 MgCl2, 1.2 NaH2PO4, 2.4 CaCl2, 25 NaHCO3 and 11 glucose (equilibrated with 95% O2 and 5% CO2, pH 7.2–7.4). For extracellular local field potential (LFP) recordings, animals were deeply anesthetized with a mixture of ketamine (100 mg/kg) and xylazine (20 mg/kg) administered intraperitoneally, supplemented with subcutaneous Lidocaine (10 mg/kg) administration at the incision site. Transcardial perfusion was performed with an ice-cold aCSF (LFP-aCSF) solution containing (in mM): NaCl (125), KCl (2.5), NaH_2_PO_4_ (1.25), NaHCO_3_ (26), MgSO_4_ (1.3), glucose (10), CaCl_2_ (2), pH 7.3-7.4. After decapitation, 350-μm horizontal sections were obtained using a Leica VT1000S vibratome in oxygenated (95% O2 and 5% CO2) ice-cold LFP-aCSF. Slices were recovered in an oxygenated LFP-aCSF at 34°C for 30 min and transferred to RT for at least 30 min until use.

### Whole-cell recordings

Whole-cell patch-clamp recordings were performed using a Multiclamp 700B amplifier (Molecular Devices) from superficial ventral CA1 pyramidal neurons located in close proximity to AAV-transduced astrocytes. Patch electrodes (4–6 MΩ) were pulled from borosilicate glass capillaries (1.5 mm OD, 1.12 mm ID; World Precision Instruments) on a Sutter P-97 puller (Sutter Instruments Company) and filled with intracellular solution containing (in mM): 140 K+-gluconate, 5 NaCl, 2 MgCl2, 10 HEPES, 0.5 EGTA, 2 ATP, 0.4 GTP, pH 7.3 (280–290 mOsm). Pipette and neuronal capacitive currents were cancelled and, after breakthrough, the series resistance was compensated and monitored. Recordings were digitized on-line and filtered at 10 kHz through a Digidata 1322A interface using Clampex 10.3 software (Molecular Devices). All experiments were designed to gather data within a stable period (i.e., at least 5 min after establishing whole-cell access).

#### Extracellular field recordings

Slices were transferred to an immersion-recording chamber superfused at 3.5 mL/min with oxygenated LFP-aCSF and visualized using infrared oblique light. GFP expression was visualized using X-Cite 110LED illumination system (XT-640-W). Schaffer collaterals were stimulated for 0.1 ms at 0.033 Hz with current ranging from 5–100 μA using a monopolar electrode (A360, stimulus isolator, WPI) placed in the stratum radiatum. The inter-pulse interval of the pair-pulse ratio measurement was 50 ms. The recording electrode was either placed in the stratum radiatum for evoked field excitatory post-synaptic potentials (fEPSPs) recording or in the pyramidal cell layer for evoked population spikes (pSpike) recording. Both stimulation and recording electrodes (1-3 MΩ) were pulled from borosilicate glass (OD 1.5 mm, ID 0.86 mm; WPI) using a horizontal flaming/brown micropipette puller (P97; Sutter Instrument) and filled with LFP-aCSF. AMPAR-mediated fEPSP and pSpike recording were performed with LFP-aCSF in the presence of 50 μM picrotoxin (HelloBio). For NMDAR-mediated fEPSP recording, slices were superfused with Mg^2+^-free LFP-aCSF (omitting MgSO_4_) in the presence of 50 μM picrotoxin and 10 μM NBQX (MedChemExpress) for at least 20 min before recording. Data were recorded with Multiclamp 700B amplifiers, filtered at 10 kHz, and sampled at 20 kHz with a Digidata 1550B interface (Molecular Devices) controlled by pClamp 10.7 software (Molecular Devices).

### Primary murine astrocyte culture

A commercial primary murine astrocyte (ScienCell, #M1800-57-SC) isolated from neonatal (P0) brain was used. Cells were previously thawed at passage 3 and were seeded at a density of 20,000 cells per P24 wells, on polylysine-coated round glass coverslips (12 mm diameter No. 1.5: Epredia Menzel, Oxford Instruments; Poly-Lysine Sigma, #P8920, 1mg/ml). Cells were maintained in 500 µl Astrocyte Medium (ScienCell, #1831-SC) for around 6 days, and culture medium was replaced every 1 to 3 days. Cells were treated for 24 h with 10 µM of PNU282987 (Bio-Techne, #2303), with or without 30 µM of AG490 (Bio-Techne, #0414) diluted in 1% DMSO (Vehicle). Cells were then washed three times for 3 min in PBS and fixed in 4% PFA for 20 min at RT.

### Histology

Mice received a lethal dose of sodium pentobarbital (180 mg/kg) and were then transcardially perfused 2 min with phosphate buffered saline (PBS) for exsanguination followed by 8 min with 4% paraformaldehyde (PFA) for fixation. Brains were post-fixed in 4% PFA for 4h and cryoprotected in 30% sucrose/0.1M PBS for 48 hours. Serial free-floating cryosections (8-10 series, 30-40 µm thickness) were obtained using a freezing microtome and stored at −20°C in a cryoprotectant solution (30% ethylene glycol, 30% glycerol, 0.1 M phosphate buffer (PB) in distilled water) until use.

#### Immunofluorescent staining on mouse brain sections

For all antibodies except STAT3α, CC1, NG2, and GS brain sections were rinsed 3 times in 0.1M PBS and blocked in 0.1M PBS/0.2%Triton X (Tx)-100/4.5% serum (normal goat or horse) for 1h at RT. Primary antibodies directed against: GFP (Aves Labs, #GFP-1020, chicken, 1:1000 or Sicgen, #AV0066-200, goat, 1:500), GFAP-Cy3 (Sigma, #C9205, mouse, 1:1000), NeuN (Millipore, #ABN91, chicken, 1:1000), td-Tomato (Sicgen, #AB8181, goat, 1:500), IBA1 (Wako, # 019-19741, rabbit, 1:1000), c-Fos (Santa Cruz, #sc-52, rabbit, 1:250-1000), vimentin (Abcam, # ab24525, rabbit, 1:1000) diluted in 0. 1M PBS/0.2%Triton X (Tx)-100/3.5% serum (normal goat or horse) were incubated overnight at 4°C under agitation. For ChAT antibodies (Merck, #AB144P, goat, 1:200 or Merck, #ab178850, rabbit, 1:500), the incubation was extended to 48-72h. The following day, sections were rinsed 3 times in 0.1 M PBS, before incubation of Specie-specific Alexa-Fluor secondary antibodies (Invitrogen, 1:500) 1h at RT in 0.1 M PBS/0.2%Tx-100/3% serum. For STAT3α antibody (Cell signaling, #8768P, rabbit, 1:500) slices were pretreated with pre-cooled absolute methanol for 20 min at −20 °C and rinsed 3 times in 0.1 M PBS before blocking and the primary antibody was diluted in SignalStain® antibody diluent (Cell signaling, #8112L) and incubated for 48-72 h at 4°C under agitation. For CC1/APC (Sigma, #OP80, mouse, 1:500), NG2 (Millipore, #ZRB5320, rabbit, 1:500) and GS (Chemicon, #MAB302, mouse, 1:1000) brain sections were rinsed in Trizma Base saline (TBS) (50 mM Trizma hydrochloride pH=7.4/0.9% NaCl), permeabilized and blocked in TBS/0.5% Tx-100/10% normal goat serum (NGS) or bovine albumin (BSA) serum for 3h at RT. Primary and secondary antibodies were incubated in TBS/0.5% Triton/3% NGS or BSA during 4 days and 2h, respectively. Slices were then incubated in DAPI (0.2 mg/ml), rinsed and mounted on SuperFrost® Plus (Thermo-Fisher Scientifc) slides and coverslipped with Fluorsave™ (Calbiochem, Darmstadt, Germany).

#### RNAscope on mouse brain sections

Sections were washed in 0.1M PBS containing 0.5%Tx-100 and 0.3% polyvinyl sulfonic acid (PVSA; RNase inhibitor; pH < 7.5). Delipidation was performed in PBS supplemented with 4% sodium dodecyl sulfate and 200 mM boric acid (pH < 7.0) for 1 h at 37°C under agitation. Following rinsing and washing steps in PBS/Tx-100/PVSA, endogenous peroxidase activity was quenched by incubation in 3% hydrogen peroxide for 10 min. After additional washes, sections were incubated with 4 drops of undiluted RNAscope™ Probe - Mm-Chrna7 (Bio-Techne, #465161) for 2 h at 40°C in a hybridization oven. Sections were then washed 3 times for 5 min in the RNAscope Wash Buffer. Signal amplification was performed using sequential incubations with AMP1, AMP2, and AMP3 reagents (30 min each, at 40°C) of the RNAscope Multiplex Fluorescent Reagit Kit (ACD biotechne, #323100), with intermediate washes in RNAscope Wash Buffer. For signal detection, sections were incubated with horseradish peroxidase (HRP) -C1 reagent for 15 min at 40°C, followed by fluorescent labeling using Opal 570 (Quanterix, #FP1488001KT, 1:1500 dilution in TSA buffer) for 30 min at 40°C. After washing, endogenous HRP activity was blocked using the HRP-blocker reagent for 15 min at 40°C. Final washes were performed in RNAscope Wash Buffer and PBS containing 0.5%Tx-100. Following RNAscope, sections were processed for immunofluorescence. Samples were first blocked in 0.1 M PBS/0.2% Tx-100/5% horse serum for 1h at RT. Sections were then incubated overnight at 4°C with a primary anti-GFP (Aves Labs, #GFP-1020, chicken, 1:1000) diluted in 0.1 M PBS/0.2% Tx-100/3.5% horse serum. After washes in PBS, sections were incubated with a donkey anti-chicken Alexa Fluor 488 secondary antibody (Invitrogen, 1:500) for 1h30 at RT under agitation. Finally, nuclear staining was performed using DAPI (1:5000 in PBS) for 20 min at RT. Sections were washed thoroughly in PBS between each step.

#### Immunofluorescent staining on primary astrocyte cultures

Following fixation, cells were washed 3 times in PBS and incubated in PBS containing 0.5% Tx-100 for 10 min for permeabilization, followed by three successive 3 min washes in PBS. Cells were then blocked by incubating for 1h in 0.1 M PBS/0.1% Tx-100/10% horse serum before overnight incubation at 4°C without agitation in primary antibody solution composed of SignalStain® antibody diluent (Cell signaling, #8112L) containing STAT3α antibody (Cell signaling, #8768P, rabbit, 1:500), PHDGH (Euromedex, #CO-NMD-MSFR100030, guinea pig, 1:400) and S100β (Synaptic System, #287004, guinea pig, 1:300). The following day, cells were washed three times for 3 min in PBS and incubated for 1h at RT in secondary antibody solution with donkey Alexa Fluor 488 secondary antibodies in 0.1 M PBS/0.1% Tx-100/10% horse serum. After PBS washes, DAPI (1:500) was incubated on cells for 10 min at RT. Finally, cells were washed three times in PBS prior to mounting in FluorSave mounting medium.

### Protein extraction and western blotting

Mice received an overdose of sodium pentobarbital and were decapitated. Brains were rapidly removed and hippocampi dissected out. 4-6 punches (1 mm) around the vCA1 area were collected between two 1-mm thick slices done on a mouse brain matrix, immediately frozen in liquid nitrogen and kept at −80°C until further processing. Tissue was thawed and homogenized in 50 mM Tris HCl (pH 7.4), 2% SDS, 150 mM NaCl, protease (1:100, Protease inhibitor cocktail, #P8340, Sigma-Aldrich) and phosphatase (1:100, Phosphatase inhibitor cocktail 2, #P5726, Sigma-Aldrich) inhibitors using a commercial homogenizer (Precellys). Protein concentration was determined by the BCA method and samples were diluted in loading buffer (NuPAGE® LDS sample buffer and sample reducing agent, Invitrogen) before denaturation at 70°C for 10 min. Samples (20 µg/well) were loaded on 4-20% Criterion™ TGX Stain-Free Precast Gels (Bio-rad) and migration was performed 1 h at 170V in Tris-glycine buffer (Bio-rad) followed by Stain-Free gel activation using a ChemiDoc MP Imaging System (Bio-rad). Proteins were transferred on a nitrocellulose membrane with the Trans-Blot Turbo™ Transfer System (Bio-Rad). Total protein load per well was evaluated after imaging the membrane under UV light. Membranes were rinsed in 9mM NaCl/Tris-HCl Buffer/0.1% Tween 20 (T-BST) and blocked in T-BST/5% BSA for 1h at RT and sequentially incubated overnight at 4°C with the following primary antibodies directed against: STAT3a (Cell signaling, #8768P, rabbit, 1:500) and GFAP (Dako, #Z0334, rabbit, 1:5000) in TBST/5% BSA. After three 10 min washes, in TBS-T, membranes were incubated for 1 h at RT with HRP-conjugated secondary antibodies (Vector laboratories, 1:5000) diluted in T-BST/5% BSA. Membranes were incubated 1 min with Clarity Western ECL substrate (Bio-rad) and the signal was detected with ChemiDoc MP Imaging System. Band intensity was quantified with Image Lab Version 5.2.1 and normalized to total loaded protein load as detected on the Stainfree membrane before antibody incubation.

### Tissue dissociation and fluorescence activated cell sorting

We used fluorescence activated cell sorting (FACS) to sort AAV-transduced astrocytes based on their expression of fluorescent proteins. Adult WT mice were bilaterally injected in the striatum with AAV-GFP + AAV-td-Tomato. One-month post-injection, mice were euthanized via cervical dislocation, their brains rapidly collected and sliced into 1mm-thick coronal sections using a brain matrix (Ted Pella). The GFP+ area was dissected on each slice using 1 mm-diameter punches under a fluorescence macroscope (Leica), pooling samples from 4 mice. Punches were collected in 1 ml ice-cold Ca2+, Mg2+ Hank’s Balanced Salt Solution (HBSS). Brain tissue was dissociated using the Neural Tissue Dissociation Kit protocol (Miltenyi Biotec, Germany). Brain punches were incubated in pre-warmed solution containing papain and DNAse for 15 min at 37°C. They were manually dissociated using fire-polished Pasteur pipettes. After several incubations/dissociation cycles, the homogenate was filtered through a 40 μm filter and diluted in 10 ml of ice-cold HBSS. Samples were then centrifuged 10 min at 300 g, the supernatant was discarded and the dissociated cells resuspended in 1 ml of HBSS. FACS was performed on an Influx biohazard cell sorter (BD Biosciences, San Jose, CA). The population of interest was determined using SSC and FSC parameters to exclude myelin debris, aggregated cells and doublets. Sample from a non-injected C57BL/6 control mouse was used to evaluate basal autofluorescence.

### Microscopy and image analysis

Images were acquired on a Leica DM6000 epifluorescent microscope (10x and 20x) or on Leica TCS SP8 confocal microscope (40x and 64x) (10-15 steps, z-step:1µm, resolution: 1024*1024px). Image analysis and histological quantifications were done using Fiji. To assess the number of transduced cells in different subregions of the hippocampus, the number of GFP+ cells were counted using ‘*Multipoint tool*’ on 20X epifluorescent tiled images. The total number of GFP+ cells per subregion was normalized to the hippocampus subregion area. The number of GFP+, STAT3+, GFAP+ cells was manually determined using the ‘*Cell Counter*’ plugin. Quantification on confocal images were performed on maximum intensity Z-stack projections. STAT3 immunoreactivity (i.e mean grey value) was measured in individually segmented astrocyte nucleo-somatic area. GFAP+ area was measured using Fiji threshold function, divided by total image area and expressed as a percentage. Three to 5 non overlapping fields of view were imaged in CA1 (*strata oriens, radiatum, lacunosum moleculare)*, CA3 and dentate gyrus, in 2-4 serial brain sections per mouse. For cell culture experiments. For data on primary astrocyte cultures, STAT3 immunoreactivity was measured nucleo-somatic area of individual cells, normalized by the averaged value obtained for the vehicle condition per culture. The mean value per well was then calculated.

### Statistical analysis

Results are expressed as mean (value per mouse) ± SEM. Statistical analysis was performed using GraphPad Prism 10 and R studio. The normality of variables/residues and homoscedasticity were assessed using Shapiro-wilk and Spearman correlation tests. Parametric tests were used if these requirements were met or when sample size was too low, as non-parametric equivalents rely on ranking, which is not reliable for small sample size. In experiments with paired samples, we used a two tailed paired t-test, a two-way repeated measure ANOVA or Mixed model with Geisser-Greenhouse’s correction followed by post-hoc tests (Sidak’s or Tukey’s multiple comparison tests). For all other experiments, we used two-tailed unpaired t-tests and one-way ANOVA followed by Tukey’s post-hoc tests for parametric data and Mann-Whitney or Kruskal-Wallis tests followed by Dunn’s post-hoc test for non-parametric data. For fiber photometry z-score traces, statistical significance was assessed using a two-sided permutation test (2,000 permutations), as described in ^50^. Linear mixed model fit by restricted maximum likelihood was performed for analyzing RNAscope and immunocytofluorescence data using the ‘lmer’ R package. A detailed description of statistical tests per panel is showed in Table 1. Statistics and sample size are described in figure captions.

